# On the role of phase lag in multi-appendage metachronal swimming of euphausiids

**DOI:** 10.1101/2020.06.30.180851

**Authors:** Mitchell P. Ford, Arvind Santhanakrishnan

## Abstract

Metachronal paddling is a common method of drag-based aquatic propulsion, in which a series of swimming appendages are oscillated, with the motion of each appendage phase-shifted relative to the neighboring appendages. Ecologically and economically important euphausiid species such as Antarctic krill (*E. superba*) swim constantly by stroking their paddling appendages (pleopods), with locomotion accounting for the bulk of their metabolic expenditure. They tailor their swimming gaits for behavioral and energetic needs by changing pleopod kinematics. The functional importance of inter-pleopod phase lag (*ϕ*) to metachronal swimming performance and wake structure is unknown. To examine this relation, we developed a geometrically and dynamically scaled robot (‘krillbot’) capable of self-propulsion. Krillbot pleopods were prescribed to mimic published kinematics of fast-forward swimming (FFW) and hovering (HOV) gaits of *E. superba*, and the Reynolds number and Strouhal number of the krillbot matched well with those calculated for freely-swimming *E. superba*. In addition to examining published kinematics with uneven *ϕ* between pleopod pairs, we modified *E. superba* kinematics to uniformly vary *ϕ* from 0% to 50% of the cycle. Swimming speed and thrust were largest for FFW with *ϕ* between 15%-25%, coincident with *ϕ* range observed in FFW gait of E. superba. In contrast to synchronous rowing (*ϕ*=0%) where distances between hinged joints of adjacent pleopods were nearly constant throughout the cycle, metachronal rowing (*ϕ* >0%) brought adjacent pleopods closer together and moved them farther apart. This factor minimized body position fluctuation and augmented metachronal swimming speed. Though swimming speed was lowest for HOV, a ventrally angled downward jet was generated that can assist with weight support during feeding. In summary, our findings show that inter-appendage phase lag can drastically alter both metachronal swimming speed and the large-scale wake structure.

## 1 Introduction

The coordinated rowing of multiple appendages is a common biological fluid transport mechanism used for diverse functions such as swimming in crustaceans [1-5], walking in echinoderms [6], ventilation in mayfly nymphs [7-9] and pulmonary mucus clearance in mammals [10-13]. Coordinated, sequential paddling of appendages generates a metachronal wave, such that adjacent members maintain a nearly constant phase difference [6]. Metachronal rowing is used by aquatic organisms spanning a wide range of sizes and swimming speeds, with Reynolds number (ratio of inertial forces to viscous forces) ranging from 10^−5^ for paramecia [14] to 10^5^ for escaping mantis shrimp [4]. A number of studies of metachronal swimming have investigated gait kinematics in relation to behavioral responses of various organisms, including paramecia [14-16], copepods [5,17-18], mysids [19], krill [20-21], mantis shrimp [4], and lobsters [22]. However, diversity of body and appendage morphologies across metachronal swimmers make it difficult to generalize how specific kinematic parameters impact swimming performance. Mechanistic studies are needed to identify unifying physical design principles underlying this successful bio-locomotion strategy. These mechanistic studies can inform evolutionary and functional biologists on the physical parameters underlying swimming performance in paddling organisms, as well as engineers working on the development of biomimetic aquatic drones.

Antarctic krill (*Euphausia superba*) are one of the most well-studied euphausiid crustaceans [1,20-21,23-32] on account of their ecological significance, with a global biomass comparable to that of humans [27]. They provide a crucial connection in oceanic food webs by grazing on smaller plankton and serving as prey for larger, commercially important animals such as fishes. Antarctic krill spend much of their lives migrating vertically and horizontally tens of kilometers each day, with locomotion costs accounting for nearly three-fourths of their daily metabolic expenditure [28, 29]. This makes minimizing cost of transport essential for their survival, rather than simply maximizing thrust or maneuverability, as would help with predation avoidance.

*E. superba* swim by periodically stroking five pairs of closely-spaced swimming appendages in an adlocomotory sequence starting from the tail to head of the animal. Motion of pleopod pairs are phase-shifted in time relative to their neighboring pairs. Jointed pleopods allow pleopod motion to be geometrically asymmetric in time by allowing the endopodite and exopodite to fold in during the recovery stroke to reduce drag. Two commonly seen swimming gaits in *E. superba* are fast-forward (FFW) and hovering (HOV) [20]. HOV is used by *E. superba* during feeding in laboratory settings [1,30], and in the wild [31]. Faster swimming speeds realized by FFW [20] can be beneficial for collective migration, coordinated schooling behavior, and for predation avoidance [32]. These gaits are defined for different swimming behaviors, but the pleopod kinematics associated with each gait exhibit significant differences in stroke amplitude (SA), phase lag between pleopods (*ϕ*), and body orientation [20]. While we know that changing gait impacts swimming speed of *E. superba* [20], the functional importance of individual kinematic parameters (specifically SA and *ϕ*) on free-swimming performance remains unclear. As changes in *E. superba* gaits involve coupled changes to two or more stroke kinematic parameters, alternative approaches are necessary to identify functional roles of individual kinematic parameters. In this regard, robotic [33] and numerical [2,34-37] models can be useful to ascertain the relative importance of morphological and kinematic parameters. Such studies can improve our understanding of organism-environmental interactions in terms of what factors (morphology, kinematics) allow particular species to fill their specific environmental niches.

Numerical studies [2,35] have shown metachronal motion to enhance thrust, while an experimental study showed that metachrony can also contribute to lift [33]. Additionally, increasing Reynolds number results in the wake travelling farther downstream [33], consistent with the observation that the wake of *E. superba* can be detected several body lengths farther downstream than the wake of smaller *E. pacifica* [25]. Assumptions inherent to numerical models have resulted in contradictory findings. For example, recent studies have suggested both that symmetric stroking about a vertical mean angle can [2,36] and cannot [37] generate forward motion at low Reynolds numbers, regardless of phase lag. Although most modeling studies have examined tethered models of paddling propulsion [33-37], Alben et al. [2] examined swimming speed for synchronous and metachronal rowing using a simple drag coefficient model. However, they did not examine the wake structure and neglected hydrodynamic interactions between the appendages and body. In summary, modeling efforts to date have not sufficiently investigated the effects of varying stroke kinematic parameters on free-swimming performance using biologically relevant body and pleopod designs.

In this study, we developed a self-propelling paddling robot that is geometrically and dynamically similar to free-swimming *E. superba*, which we used to examine how changing inter-pleopod phase lag (*ϕ*) impacts swimming speed and forward force (thrust) generation. This model, referred to as “krillbot”, is used to examine the swimming performance and the wake structure of the HOV and FFW kinematics when changing *ϕ*. We also consider changes in body orientation angle to examine whether the larger inclination used during HOV of *E. superba* can assist with weight support by directing the wake jet more downward. Finally, we examine the time-variation in distance between adjacent pleopods in order to identify the physical mechanism underlying the superior performance of metachronal rowing compared to synchronous rowing.

## 2 Methods

### 2.1 Geometric scaling

We developed a programmable robot to model metachronal swimming with five pairs of paddling appendages. For geometric scaling of the krillbot body, morphological measurements were acquired on a high-resolution image of an Antarctic krill [38] (**Figure 1A**). These measurements were used to develop simplified geometries for the krillbot body by scaling dimensions in terms of pleopod length (**Figure 1B**). Length of crustacean pleopods varies along the length of the body, but was maintained constant in the krillbot. In krill, a joint approximately halfway down each pleopod joins the protopodite (upper portion of the pleopod) with the endopodite and exopodite (two lobed structures that make up the lower portion of the pleopod). This allows the pleopod to unfold during power stroke (PS) and fold during recovery stroke (RS), creating a drag asymmetry that helps generate forward motion. Pleopod joints were modeled using 3.2 mm diameter stainless steel rods. Pleopod shape (physical model) was simplified to a rectangular planform protopodite, and endopodite and exopodite were combined into a trapezoidal flat plate (**Figure 1C**). The krillbot pleopods were positioned at an angle of 10° from vertical, similar to *E. superba*. The front edge was squared, while the back edge was rounded, allowing the lower portion of the limb to rotate freely during the recovery stroke. This allowed the hinge to passively follow hydrodynamic forces generated by the paddling motion, but limited to a maximum *β* of 180^°^, but with no lower limit, as it is unknown whether crustaceans actively control their joints or allow them to passively follow the fluid. The tail and body angles were designed to approximately match those seen in hovering krill [20]. The body angles of 0° and 20^°^ were chosen as test conditions since 0^°^ is the minimum body drag condition for forward swimming, while 20^°^ is within the range of body angles reported for krill in both the HOV and FFW gaits [20].

**Figure 1.**
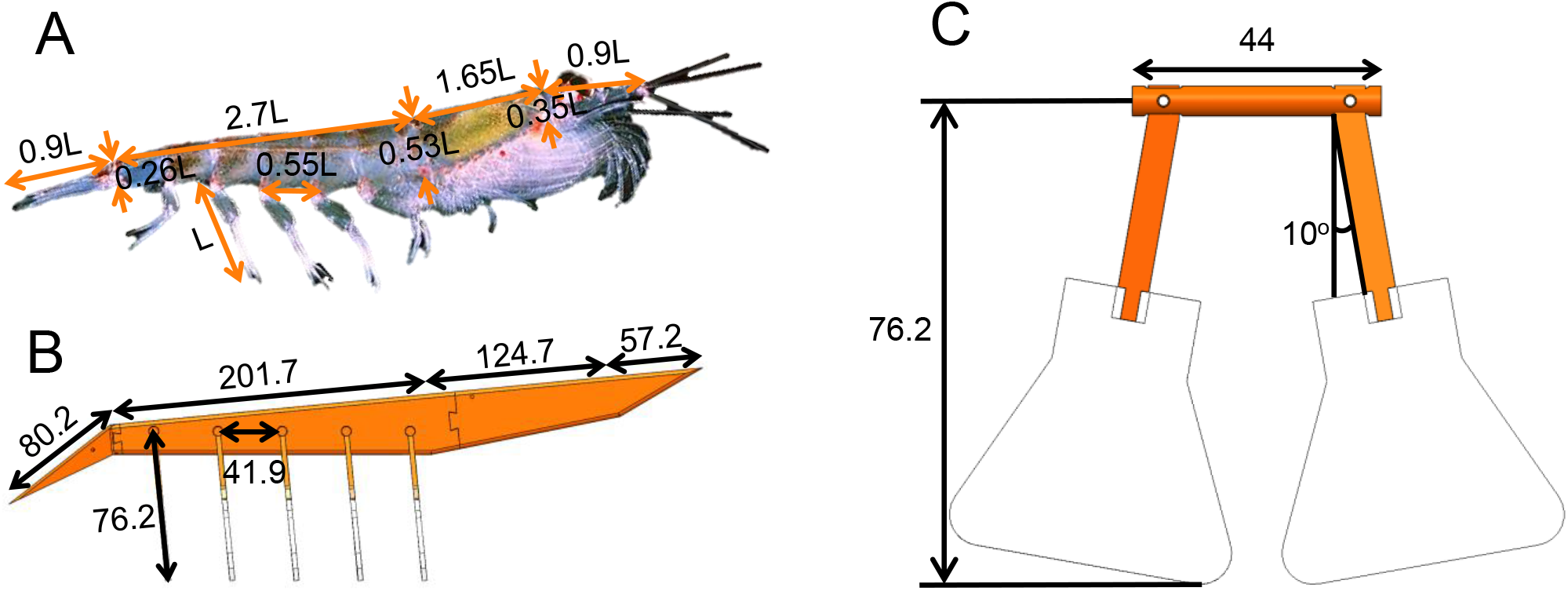
Geometric scaling of krill and krillbot design. (A) Antarctic krill (*Euphausia superba*) with lengths nondimensionalized relative to the pleopod length (L). Image of *E. superba* adapted and edited from Kils [37]. (B) Dimensions of the krillbot specified in mm. The body length of the robotic model is approximately 10 times that of adult *E. superba* and is geometrically similar. (C) Front view of a pair of krillbot pleopods (dimensions in mm). Geometry is simplified such that all pleopods are of the same length and a hinge is located midway down the length of each pleopod. Below the hinge, pleopod geometry is simplified to a trapezoidal flat plate. Rigid, solid pleopods are used on the krillbot as compared to the pleopods of *E. superba* with long feather-like setae at the fringes.

### 2.2 System design and kinematics

Motion of each pair of pleopods on the krillbot was controlled by a NEMA-23 stepper motor (model ST23-4, National Instruments Corporation, Austin, TX, USA). Pleopod motion was driven by 6.4 mm timing belts that allowed the krillbot to be submerged below the fluid surface, while motors were positioned 100 mm above the fluid surface. A custom LabView (National Instruments Corporation, Austin, TX, USA) program prescribed angular positions to each stepper motor at 10 ms increments, allowing independent control of each pleopod. The LabView program communicated with the stepper motors through a National Instruments compact RIO system (cRIO-9066) connected to five stepper motor drives (SMD-7611, National Instruments Corporation) that were used to micro-step the motors to an angular resolution of 20,000 steps per revolution for precise angular control. Experiments were performed in a 2.43 m long glass aquarium, measuring 0.65 m in width and 0.77 m in height (A). The krillbot was suspended from a 1 m long low-friction air bearing (model A-108.1000, PI (Physik Instrumente) L.P., Auburn, MA, USA) that was mounted to a custom aluminum frame built around the aquarium. A pair of stainless steel D-shafts suspended the krillbot from an aluminum frame, which was mounted either directly to the air bearing (for 0° body angle) or to a 20° wedge that was fixed to the bottom surface of the air bearing (for 20° body angle). The air bearing allowed the krillbot to move freely along the horizontal axis, driven by the hydrodynamic forces generated from the paddling motion. Filtered air was supplied to the air bearing at 550 kPa to remove frictional effects on the suspension mechanism. The primary losses (additional to hydrodynamic drag on the body) came from fluid drag on the timing belts, and from friction on the power cables used to control the motors.

Time-variation of pleopod root angle (*α* in **Figure 2D**) was prescribed to a stepper motor to drive the upper part of each pleopod pair, based on individual pleopod kinematics reported for *E. superba* performing FFW and HOV kinematics [20]. To achieve different swimming performance for different behaviors, *E. superba* change the kinematics of limb motion, as well as the angles of their body and tail relative to the direction of the flow. *E. superba* paddle their pleopods with non-uniform *ϕ* between adjacent pairs. **Table 1** shows the experimental conditions and various diagnostic measurements used in this study. For this study we tested kinematics with the unmodified, non-uniform phase lags (NU in **Table 1**) and kinematics modified to have uniform *ϕ* of 0%, 15%, 25% and 35% of the cycle (for both FFW and HOV), and 50% of the cycle (for HOV). 50% phase lag was not achievable using FFW kinematics because the large stroke amplitude and phase lag caused adjacent paddles to collide and break. Additionally, experiments for each condition were conducted with body angles of 0° and 20°. For both unmodified and modified *E. superba* kinematics, amplitude variation of *α* (indicative of SA) for each pleopod pair was prescribed to be identical to that of the equivalent pleopod pair of *E. superba*. For modified *E. superba* kinematics, only the phase-shift between adjacent pleopods was altered to be equal (varying from 0% to 50% as noted earlier). Stroke frequency of each pleopod was identically maintained at 2.5 Hz, and non-dimensional time *τ* was defined based on the cycle time *T* (i.e., *τ* = *t/T*; *T*=1 cycle=0.4 s). Start time (*t*= 0 s) was defined to be the start of power stroke of the most posterior (P5) pleopod, so that *τ*=0, 1, 2 … coincide with the start of consecutive power strokes of P5. Pleopod root angle (*α*), and hinge angle (*β*, **Figure 2D**) formed at the joint where the upper and lower parts of a pleopod meet, were tracked from high speed videos (HSVs) using ImageJ [39].

**Table 1.**
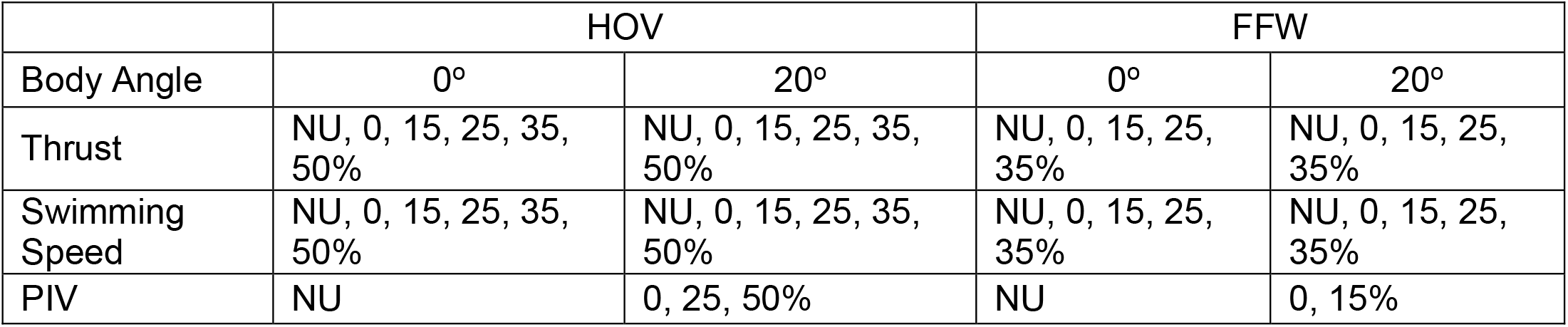
Experimental conditions used in this study. For each condition (HOV or FFW kinematics, 0° or 20° body angle), and each data type (thrust, swimming speed, and PIV), the prescribed phase lags presented in this study are shown. NU represents the unmodified animal kinematics, while numeric values represent uniform values of *ϕ*, as a percentage of cycle time.

**Figure 2.**
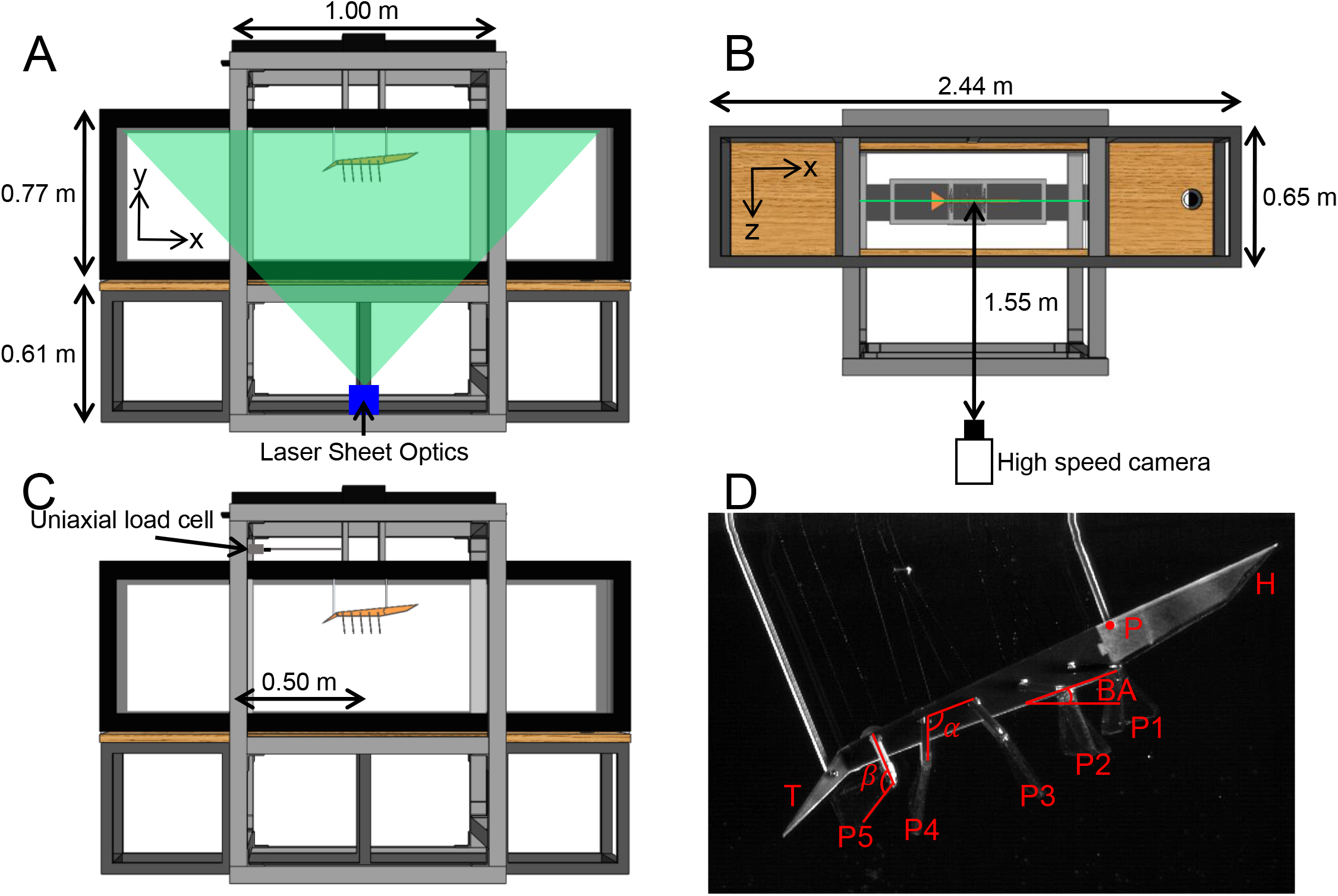
Experimental apparatus. (A) Camera view used in PIV data acquisition. A laser sheet illuminates seeding particles in the tank from below. The krillbot is suspended from a 1m long linear air bearing and can move forward or backward by paddling a series of pleopods. (B) Top view of the PIV setup, showing the location of the PIV laser plane. Clear acrylic pleopods allow laser light to pass through. (C) Schematic of the experimental setup used for force measurements, made using a uniaxial load cell. (D) Close-up image of the krillbot from a representative high-speed video used to quantify swimming speed. Position of the mounting screw (P) near the head (H) of the model was tracked in each frame. Pleopods are labeled from anterior (P1) to posterior (P5). Time-variation of pleopod root angle (*α*) was prescribed, and pleopod joint angles (*β*) were allowed to passively fold during recovery stroke (RS) and unfold in power stroke (PS).

Tracking was performed at a reduced time-resolution of 25 frames/second (every 8th frame of HSVs acquired at 200 frames/second). Since the pleopods were mounted at an angle relative to the body, this resulted in some distortion of the *β* angle and introduced some uncertainty due to the out-of-plane rotation of the hinge. Additionally, since the hinges of each pleopod operated independently, it was possible for the left and right pleopods in a pair to have different *β* values at any given time. The results of the kinematics tracking for the pleopod closer to the camera are shown over the last full cycle of each trial (**Figure 3**), to ensure that the model had reached a steady swimming speed. Mean and standard deviations were obtained across five independent trials. Representative HSVs are provided in electronic supplementary material.

**Figure 3.**
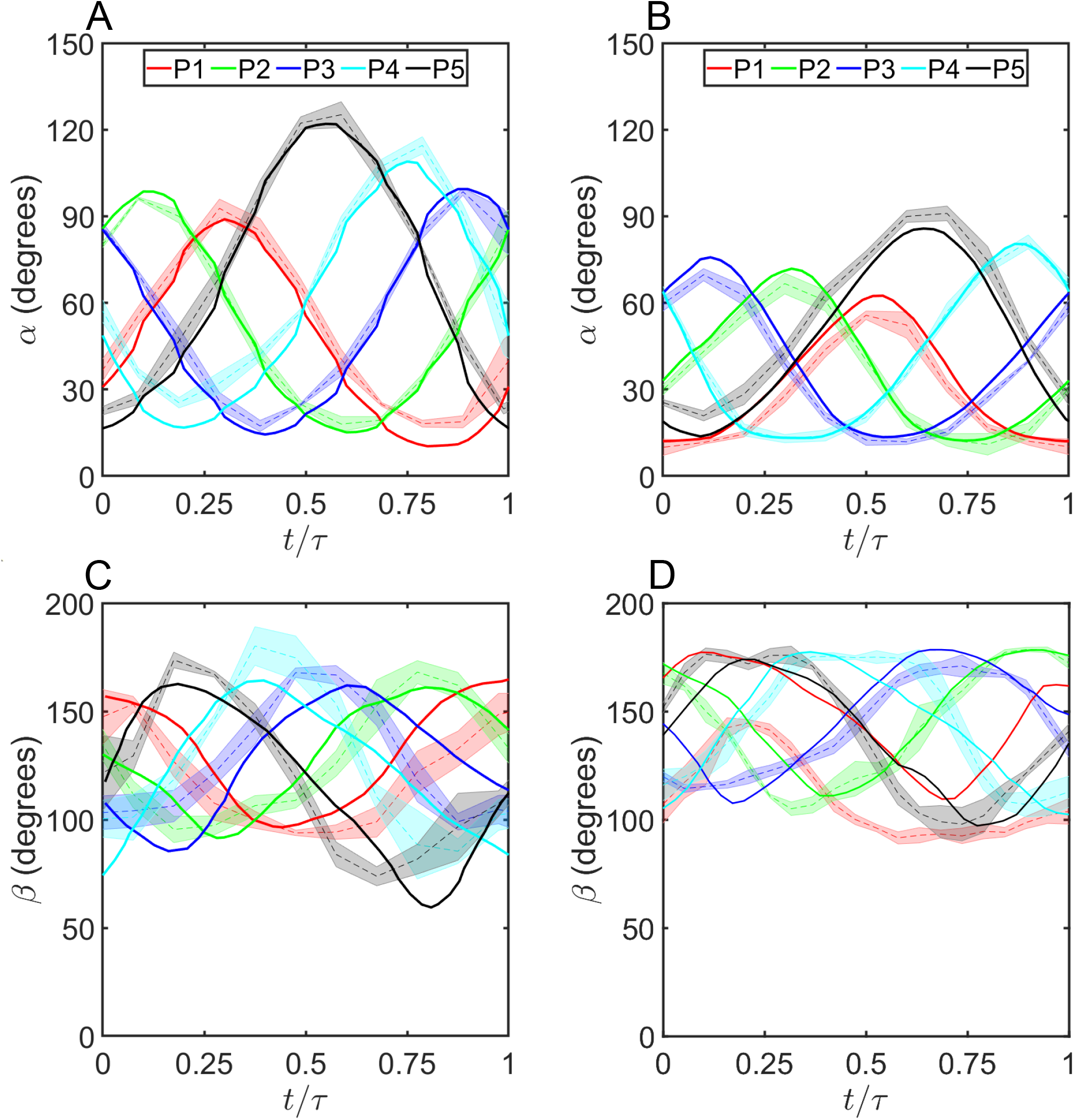
Pleopod kinematics of Antarctic krill [20] and of the krillbot. (A) Pleopod root angle (α) versus time for non-uniform (NU, **Table 2**) fast-forward (FFW) kinematics. Solid lines represent prescribed motion, obtained from *E. superba* [20] and dashed lines represent the mean kinematics achieved by the krillbot (shading represents ±1 standard deviation across 5 trials). (B) *α* versus time for non-uniform (NU, **Table 2**) hovering (HOV) kinematics. (C) Hinge angle *β* versus time for FFW kinematics. Hinge angle *β* was not actively controlled, and followed the hydrodynamic forces generated by the paddling motion. (D) *β* versus time for HOV kinematics.

### 2.3 Dynamic scaling

The test aquarium was filled with a mixture of 85% glycerin and 15% water by volume (density, *ρ*=1225 kg/m^3^; kinematic viscosity, *ν*=100 mm^2^/s). This mixture allowed for matching the Reynolds number (*Re*) of flow generated by krillbot to that of freely-swimming Antarctic krill, for dynamic similarity. Pleopod *Re* was defined as:

**Table 2.** Metrics used to compare the krillbot swimming performance to the swimming performance of live *Euphausia pacifica* and *Euphausia superba*. The kinematics used in this study came from *E. superba* [20]. Data for *E. pacifica* and *E. superba* obtained from [20,25]. Displacement efficiency, advance ratio, and Strouhal number allow for direct comparison to the live animal swimming performance. Displacement efficiency is lower in the krillbot than in krill, resulting in a lower advance ratio.

| Metric | Swimming mode | *E. pacifica* | *E. superba* | Krillbot |
| --- | --- | --- | --- | --- |
| Body Length (cm) | N/A | 2.7 | 1.0-5.5 | 46.4 |
| Limb Length (cm) | N/A | 0.25 | 0.65 | 7.62 |
| Number of Pleopod Pairs | N/A | 5 | 5 | 5 |
| Maximum Stroke Amplitude (°) | HOV | — | 76 | 71 |
|  | FFW | 111 | 105 | 102 |
| Pleopod Beat Frequency (Hz) | HOV | 5.6 | 3 | 2.5 |
|  | FFW | 10.9 | 6.2 | 2.5 |
| Swimming Speed (cm/s) | HOV | 2.6 | 1 | 4.9 |
|  | FFW | 15.5 | 12.6 | 15.3 |
| Normalized speed (BL/s) | HOV | 1 | 0.26 | 0.1 |
|  | FFW | 4.84 | 4.0 | 0.33 |
| Displacement Efficiency (BL/stroke) | HOV | 0.46 | 0.09 | 0.04 |
|  | FFW | 0.44 | 0.65 | 0.13 |
| Advance ratio | HOV | 0.096 | 0.039 | 0.02 |
|  | FFW | 0.29 | 0.19 | 0.045 |
| Reynolds Number | HOV | 135* | 310-360 | 360 |
|  | FFW | 264 | 380-1400 | 517 |
| Strouhal Number | HOV | — | 0.37 | 0.44 |
|  | FFW | — | 0.38 | 0.45 |

\* = stroke amplitude for FFW *E. pacifica* used for calculations

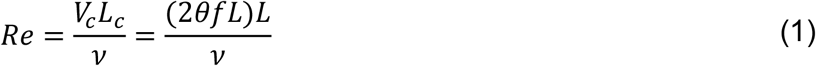

where *V*_*c*_ and *L*_*c*_ are characteristic velocity and length scales, respectively. We chose pleopod length (*L*) as *L*_*c*_ and the mean pleopod tip speed for *V*_*c*_. *V*_*c*_ is the product of stroke frequency (*f*) and arc length (2*θL*). *θ* represents SA of the tail-most pleopod (SA is non-uniform among pleopods [20]). *Re* of krill during HOV gait ranges from about 310-360, while *Re* of the krillbot performing HOV kinematics is 360 (**Table 2**). For the fluid dynamic characteristics of the flow to match, both the Reynolds number and the Strouhal number should be matched. The Strouhal number (*St*) is defined as follows:

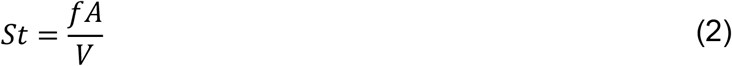

where *f* is the dominant frequency in the flow (equivalent to the stroke frequency), while *A* and *V* are the characteristic length and velocity, respectively. Since paddling crustaceans are often found hovering, Murphy et al. [21] proposed using the maximum velocity in the wake (*V*_*wake*_) as the characteristic velocity for swimming speeds less than 2 body lengths per second. For the characteristic length, they used the horizontal distance traveled by the pleopod tip during PS (*A* = 2*L sin*(*θ* / 2)). These definitions are used here for comparison. Based on this definition, Strouhal number is much less sensitive to changes in limb kinematics than Reynolds number. If it is assumed that the velocity in the core of the wake is approximately equal to the mean tip speed of the paddles (2*θfL*), then the Strouhal number ranges from 0.5 for *θ*=0° to 0.32 for *θ*=180°, and is dependent only on *θ* (consistent with the *St* range reported for *E. superba* [21]). This also overlaps with the range of *St* associated with the most efficient swimming performance in a number of other flying and swimming organisms [40]. *Re*, on the other hand, is linearly dependent on both the stroke amplitude and stroke frequency, as well as the square of the paddle length. Since both *Re* and *St* values achieved by the krillbot are similar to those of the krill (**Table 2**), physical characteristics of the flows generated by the krillbot and by *E. superba* are expected to be comparable for similar normalized swimming speeds.

### 2.4 Swimming speed

Krillbot swimming speed was determined from high-speed videos (HSVs) acquired at 200 frames/second using a high-speed camera (Phantom Miro M110, Vision Research, Wayne, NJ, USA). A 50 mm constant focal length lens (Nikon Micro Nikkor, Nikon Corporation, Tokyo, Japan) was attached to the camera (aperture setting of f/2.8) for all HSV measurements. The camera was placed 2.3 m from the mid-plane of the krillbot body to give a 1.2 m long FOV, to permit recording the entire length of krillbot travel (along the 1 m long air bearing). Displacement of a fixed point (P in **Figure 2D**) was tracked in time using DLTdv7 [41] in MATLAB (The MathWorks, Inc., Natick, MA, USA). Displacement of the krillbot was tracked in time, with averages and standard deviations taken over five independent trials. Displacement for the prescribed animal kinematics is shown in **Figure 4A**. A higher slope of the displacement curve means that the krillbot moves faster. Body velocity was defined as the slope of the displacement curve, but was found to vary greatly within a cycle (**Supplementary Figures S1-S3)**. Because of this, we defined a mean swimming speed (*V*_*mean*_) to characterize swimming performance:

**Figure 4.**
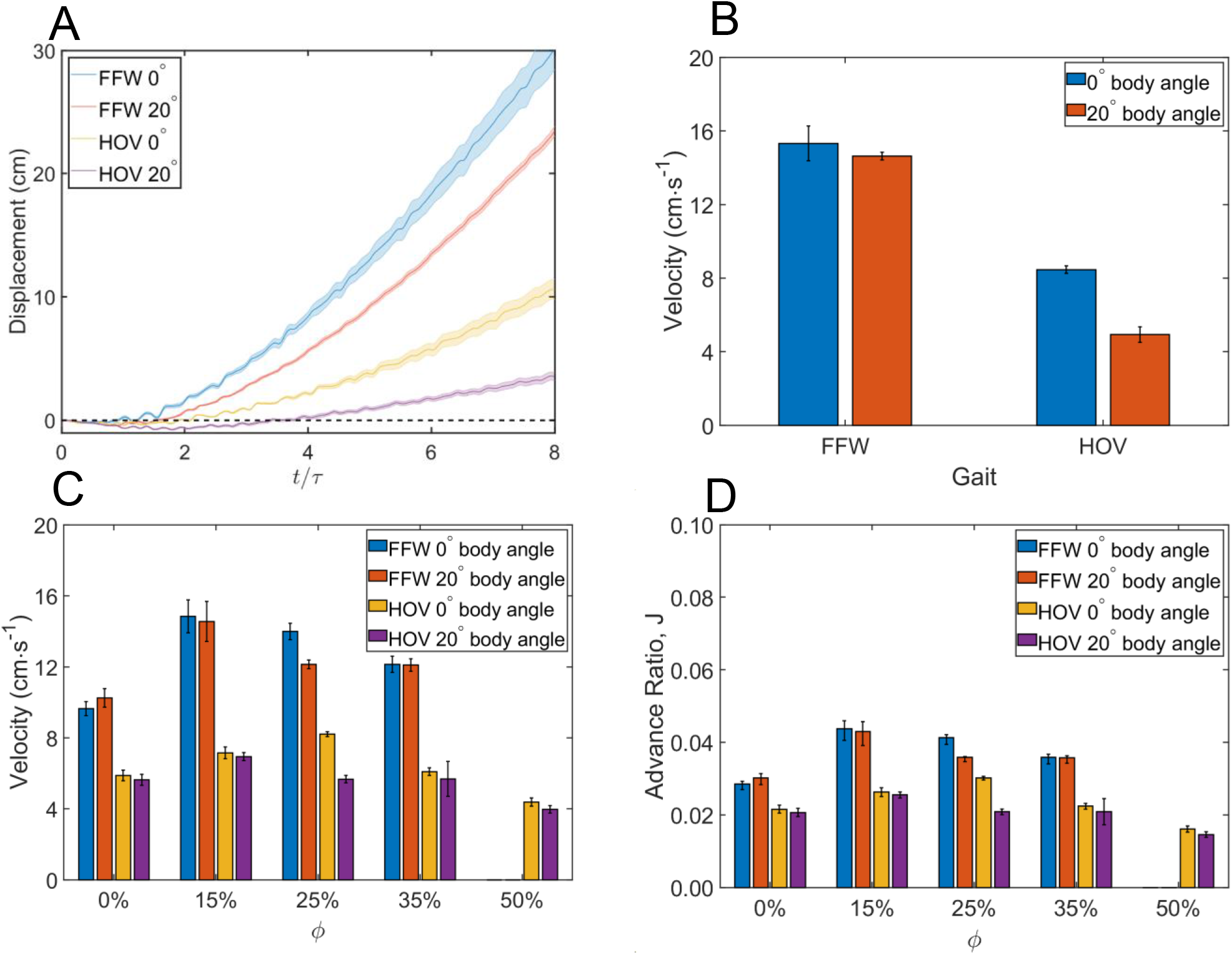
Self-propulsion characteristics of the krillbot. (A) Krillbot displacement versus non-dimensional time (*τ*) for non-uniform *ϕ*. (B) Steady swimming speed for the conditions shown in (A). (C) Steady swimming speed for varying phase lag and kinematics for modified *E. superba* kinematics with uniform *ϕ*. For both FFW and HOV, uniform 15% phase lag shows nearly equal swimming speed (in C) as non-uniform phase lag (in B). (D) Advance ratio for varying kinematics, body angle, and phase lag, calculated using equation (6).

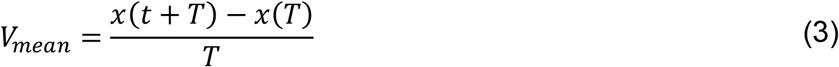

where *x*(*t*) is displacement at time *t, T* is cycle time, and *x*(*t* + *T*) is displacement in one cycle after time *t*. This definition is the 1st order forward derivative in discrete time for one cycle. Mean swimming speed increased over the first several cycles, but achieved a constant value (asymptotic slope in **Figure 4A**) that was recorded as *V*_*mean*_. Mean and standard deviations of *V*_*mean*_ were calculated across five independent trials. Representative HSVs are provided in electronic supplementary material.

### 2.5 Thrust

An advantage to using a robotic model is that the krillbot allows for force measurements in a repeatable and controllable manner that may not be possible in organismal studies. In this study, thrust measurements were acquired for different phase lags for FFW and HOV kinematics at body angles of 0° and 20°. Forward propulsive force (thrust) was recorded using a 250 g uniaxial load cell (GSO-250, Transducer Techniques, Temecula, CA, USA). One end of the load cell was attached to the rear suspension rod between the krillbot and air bearing (**Figure 2C**), while the other end was connected to the aluminum frame via a short tether. Time-averaged thrust was calculated over each cycle, and the mean and standard deviation of time-averaged thrust were taken over 20 cycles (after forces reached a steady state). Time-averaged thrust 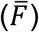 was defined as:

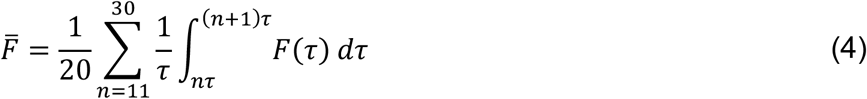

where *n* is the cycle number, *F* is the time-varying thrust measured by the load cell, and *τ* is dimensionless time (*τ* = *t/T*; *T*=1 cycle=0.4 s). Thrust measurements were acquired for *E. superba* FFW and HOV kinematics [20] and for modified *E. superba* kinematics with *ϕ* ranging from 0% to 50%, at body angles of 0° and 20°. It is important to note that the FFW and HOV swimming kinematics may not be the best-suited for accelerating from rest, as tethered organisms have been observed to greatly change their behaviors when paddling [17,42].

### 2.6 Particle image velocimetry (PIV)

PIV measurements were conducted in order to visualize the flow field caused by the paddling motion and the vorticity in the wake. The velocity fields show direction and magnitude of the flow at discrete points in the wake, and permit us to examine the wake structure, which is not achievable in bulk measurements such as swimming speed and thrust. PIV was performed in the mid-plane of the pleopod closest to the camera, since the pleopods of both krill and the krillbot flare out from the body, with a small space between pleopods in a pair. This allowed us to examine flow near the pleopod, where it should be the fastest and strongest. Two-dimensional, two-component PIV measurements were conducted in single-frame mode at 50 frames per second using an sCMOS camera (Imager sCMOS, LaVision GmbH, Göttingen, Germany). The fluid in the tank was uniformly seeded with titanium dioxide filled polyamide particles (55 μm mean diameter). Illumination was provided using a 527 nm wavelength high-speed laser with maximum repetition rate of 10 kHz and pulse energy of 30 mJ (Photonics Industries International, Ronkonkoma, NY, USA). The laser beam was passed through collimating optics (diverging lens and converging lens) and rotated 90° using a high-reflectivity mirror, after which it was passed through a cylindrical lens (−10 mm focal length) to generate a planar sheet (A). The camera acquired images at 50 frames/second, with image resolution of 2560 x 2160 pixels (pixel size: 6.5 x 6.5 microns). A 50 mm constant focal length lens (Nikon Micro Nikkor, Nikon Corporation, Tokyo, Japan) was attached to the sCMOS camera with an aperture setting of f/2.8 for all PIV measurements. The front of the lens was positioned 1.9 m from the mid-plane of krillbot body (lateral view), providing a field of view (FOV) of 0.63 m (length) x 0.53 m (height), with spatial resolution of 242 μm/pixel (**Figures 2A-2B**).

PIV images were acquired when the krillbot passed through the FOV, which was located near the end of the air bearing to allow enough time for the krillbot to accelerate to a steady swimming speed. PIV data were acquired for each phase lag and kinematic condition at 20° body angle. Velocity fields were calculated by multi-pass cross-correlation of raw images in DaVis 8.4 (LaVision GmbH, Göttingen, Germany). One pass each with window size of 64×64 pixels and 32×32 pixels, each with 50% overlap, was used for cross-correlation. Post-processing was performed to remove velocity vectors with peak ratio Q < 1.2. The body of the krillbot was manually traced for masking, and the masked region was excluded from the PIV cross-correlation calculations.

To examine rotational motion in the flow fields, out-of-plane (*z*) component of vorticity (*ω*_*z*_) was calculated using the following equation:

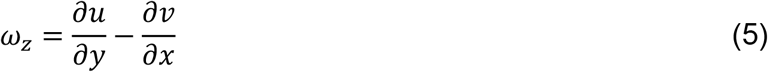

where x and y represent horizontal and vertical coordinates, respectively; and *u* and *ν* represent horizontal (*x*-component) and vertical (*y*-component) velocity, respectively. Wake vorticity has been tied to hydrodynamic signaling between neighboring individuals in aggregations [25].

### 2.7 Performance Metrics

The advance ratio (*J*), is a common measure of propulsive efficiency for forward motion that compares mean swimming speed to mean propulsor speed. While it is typically used for single propulsors rotating at a constant rate, it has been modified for use in crustaceans to account for the number of pleopod pairs (*m*) as shown below [21]:

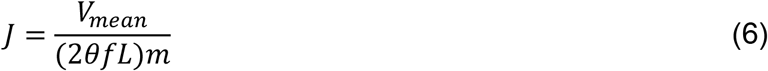

A high value of *J* means that the animal moves faster than its appendage. For drag-based propulsion (where the pleopod moves back and forth in the direction of animal motion), it would be unrealistic to have a value of *J* greater than 1. To get a value less than 1, Murphy [21] introduced a factor of 1/*m* to the advance ratio calculation. This is the definition of the advance ratio used here, so that the krillbot data can be compared directly to the data reported for *E. superba*.

As a way to determine whether the krillbot gets a boost in speed when limbs move closer together forcing a jet to escape between paddles to “push” the krillbot forward, or when the limbs move apart, resulting in the formation of strong tip vortices creating suction on the front edge of the paddles to “pull” the krillbot forward, we calculated the time-varying distance between the joints of adjacent pleopods. This was compared to the variation in swimming speed within the paddling cycle to determine when in the cycle the krillbot was moving faster or slower than average. Inter-hinge distances were calculated as:

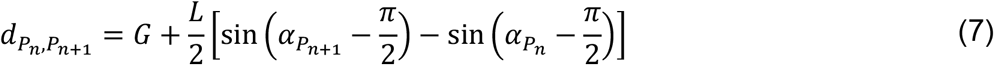

where G is the gap between pleopods (41.9 mm), 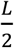 is the distance from the pleopod root to the hinge (38.1 mm), and *α* is the pleopod root angle in radians. The pleopod number n is allowed to range from 1 to 4, so when n=1 is selected, equation 7 gives the distance between the hinge joints of P1 and P2 based on the gap between limbs and instantaneous values of *α* for P1 and P2. Distances were based on manually tracked values of alpha, similar to those shown in Figures 3A and 3B.

Instantaneous deviation of the body from the mean swimming speed was calculated as the difference of the actual body position in time and the estimated body position when moving at the mean velocity, as below:

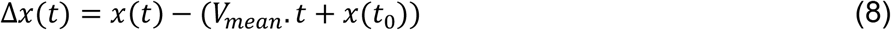

where *Δx*(*t*) is the positional deviation at time *t, x*(*t*) is the position of the robot at time *t, V*_*mean*_ *· t* is the estimated position when moving at constant speed *V*_*mean*_, and *x*(*t*_0_) is the displacement at some time *t*_0_ after the krillbot has reached a periodic steady speed. This was used as the indicator of the aforementioned speed boost that could be used to identify the mechanism of force augmentation prevalent in metachronal swimming. An increasing value of *Δx* (positive slope) will indicate a speed boost, while a decreasing value of *Δx* (negative slope) would indicate slowing of krillbot motion.

## 3 Results

### 3.1 Kinematics

For both FFW and HOV kinematics (**Figure 3**), measured pleopod root angles (dashed lines) match well with the prescribed animal data (solid lines), meaning that our model replicated the animal kinematics reasonably well. The hinge angles (*β*) are not controlled and are driven by hydrodynamic forces acting upon the pleopods. For the most part, hinge angles are qualitatively similar to those of *E. superba* [20], with the notable exception of P1 during HOV, which often failed to unfold fully during the power stroke, and had a lower average value than in the animal. Since the hinge angles are not directly controlled, it is unsurprising that they do not match as well as the pleopod root angles. *β* does not unfold as much in FFW as in HOV kinematics. *β* agrees better during FFW than during HOV. During FFW, the motion agrees relatively well in time, but with the largest differences occurring at the peak and minimum values of *β*. However, during HOV the most difference between tracked krillbot data versus krill data is in time (with the notable exception of P1). In the krill, the limb joint only briefly maintains the maximum and minimum *β* values, while the krillbot spends most of the power stroke with the hinge angle at approximately 180° *β*. A number of factors could contribute to differences between animal and krillbot *β*, including differences in energy dissipation (friction), energy storage (“springiness” of the joint), and angle limits based on the structure of the joint.

### 3.2 Swimming Speed

For both body angles (0° and 20°), FFW outperforms HOV by travelling the full length of the air bearing faster. Although the krillbot starts from rest and initially accelerates, it reaches a steady swimming speed after 7-8 stroke cycles. Using the definition of mean swimming speed from equation 3 (slope of the displacement curve in **Figure 4A**), we calculated the mean and standard deviation for each test condition (**Figure 4B**) for *E. superba* kinematics taken from Murphy et al. [20], at body angles of 0° and 20°. These are compared to mean swimming speed for modified *E*. superba kinematics with uniform phase lags of 0%, 15%, 25%, 35%, and 50% of the cycle (**Figure 4C**). We note that 50% phase lag was unachievable for FFW kinematics because the large stroke amplitude caused the paddles to collide and break. Steady swimming speed of the 15% FFW case is the same as that obtained using the non-uniform FFW kinematics of *E. superba*. The steady swimming speed using HOV kinematics of *E. superba* (non-uniform *ϕ*) is similar to that obtained using modified *E. superba* HOV kinematics with uniform *ϕ*=25%. For both FFW and HOV, maximum swimming speed occurs in the range of 15% to 25% phase lag, which agrees with expected results based on a number of previous studies on robotic (tethered) and numerical models [33,35-36]. The advance ratio (Equation 6) relates the swimming speed to the speed of the paddle and is shown in **Figure 4D**. A low value of the advance ratio means that more energy is expended in trying to move forward over the same distance. The lower advance ratio seen during HOV was expected, as most of the energy exerted by the hovering krill should go to supporting weight, rather than propelling the body forward. The lower ratio of *J*_*FFW*_/*J*_*HOV*_ as compared to *V*_*FFW*_/*V*_*HOV*_ is consistent with the expectation of increasing body drag at higher swimming speeds.

### 3.3 Thrust

The krillbot allowed for the collection of thrust data in a repeatable manner with controllable kinematics, which is not possible in organismal studies. Average thrust generated by the motion of the paddling limbs is shown in **Figure 5A** for when the krillbot is tethered and performing the *E. superba* kinematics with non-uniform *ϕ*. FFW kinematics generate more thrust than HOV kinematics, regardless of whether the body is oriented at 0° or 20°. **Figure 5B** shows the effect of changing uniform phase lag (modified *E. superba* kinematics) on time-averaged thrust for each kinematic condition and body angle. Regardless of phase lag, FFW kinematics generated more thrust than HOV kinematics. The highest thrust produced by FFW kinematics was obtained when using the animal kinematics, 1.98 ± 0.05 *N* for 0° body angle and 1.55 ± 0.05 *N* at 20° body angle. Uniform *ϕ* =15% performed most like the unmodified FFW kinematics, with 1.89 ± 0.02 *N* at 0° body angle and 1.44 ± 0.05 *N* at 20° body angle. *ϕ* =0% resulted in the least thrust for FFW kinematics, and *ϕ* =50% was unachievable during FFW motion due to the large stroke amplitude causing collisions between neighboring paddles.

**Figure 5.**
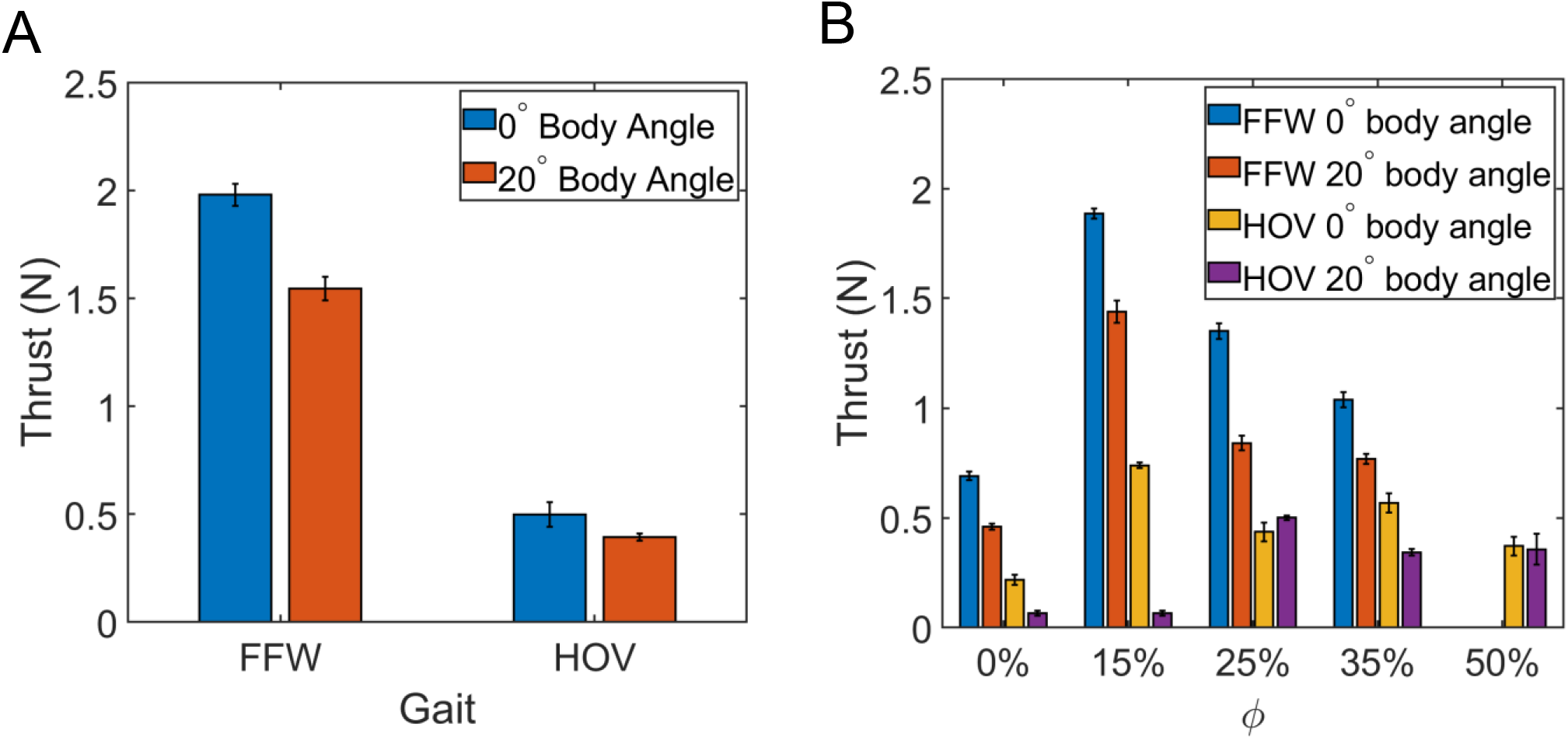
Time-averaged thrust (forward force) generated for varying *ϕ* and body angle, measured on a uniaxial load cell (**Figure 2C**). Regardless of phase lag and body angle tested, FFW kinematics (both unmodified *E. superba* kinematics and modified *E. superba* FFW kinematics with uniform *ϕ*) generated more forward force than HOV kinematics. For FFW, non-uniform phase lag (in A) was found to generate nearly the same thrust as uniform (15%) phase lag (in B).

Unlike unmodified FFW kinematics, the unmodified HOV kinematics did not result in higher thrust measurements than the modified kinematics with uniform *ϕ*. Unmodified HOV kinematics resulted in 0.50 ± 0.06 *N* thrust at 0° body angle, which is similar to the 0.44 ± 0.04 *N* measured with uniform *ϕ* =25% for the same body angle, but much lower than the 0.73 ± 0.01 *N* measured at a uniform *ϕ* =15%. Since 0° body angle is more appropriate for forward swimming and is not commonly seen in hovering krill [20], the 20° body angle case may be a more realistic case for determining the forces on hovering krill. While *ϕ* =0% and *ϕ* =15% show very almost no thrust, *ϕ* =25% has the highest thrust of 0.50 ± 0.01 *N*, which is larger than the thrust in the unmodified HOV kinematics at 0.39 ± 0.02 *N*. While the lower thrust produced with the unmodified HOV kinematics than with the uniform 25% phase lag may not seem beneficial for swimming performance, for hovering the force vector should be directed in a primarily downward direction (producing more lift than thrust). Since the direction of the wake jet cannot be determined from bulk measurements such as the forward swimming speed and thrust, PIV was used to visualize the structure of the wake.

### 3.4 Flow visualization

PIV was used to visualize the flow generated by the paddling motion under different kinematic conditions for the different body angles examined in this study. Hovering Antarctic krill have been reported to have body angles ranging from approximately 10° to 60° and phase lags ranging from approximately 15% to 30% of cycle time [20]. Since the Reynolds number, Strouhal number, and nondimensional swimming speed of the krillbot performing HOV kinematics match well with those of the krill performing the same stroke, the dynamics of the flow field should also be well matched. **Figure 6** shows the flow field generated by the krillbot performing HOV kinematics at body angles of 0° and 20°, at time points 0%, 25%, 50%, and 75% of cycle time as defined by the tail-most pleopod P5. These time points correspond to the start, middle, and end of PS and to the middle of RS. End PS also corresponds to the start of RS, while end RS corresponds to start PS. The flow field at 0° is directed in a primarily ventral (horizontal) direction, with distinct jets coming from the pleopods and from the flow past the tail. At 20° body angle, the primary jet generated by the pleopods is directed in a more downward direction, as should be expected due to the larger body angle. However, the wake is not a simple rotation of the flow generated by the HOV motion at 0° body angle. The primary jet generated by the pleopods is stronger and has clearly defined shear layers at its leading edge. The wake past the tail is also rotated in this case. The direction of the wake is directly correlated to the direction of the thrust vector, and an organism swimming in a quiescent fluid relies on the acceleration of fluid for force generation. A continuous wake directed in a primarily downward direction would help a hovering organism to maintain a constant position in the water column, rather than having to continuously adapt kinematics to move up or down.

**Figure 6.**
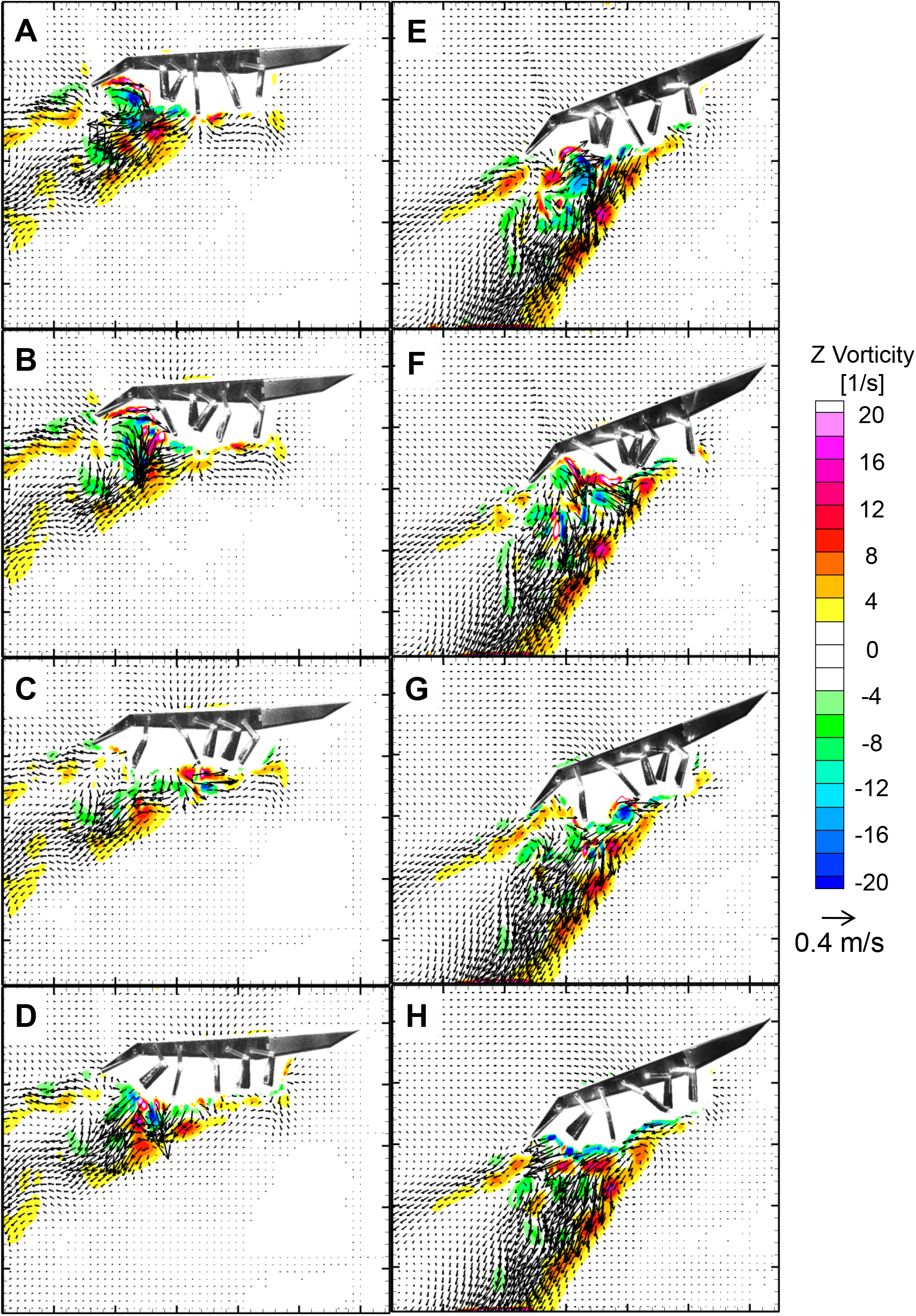
Out-of-plane vorticity contours overlaid with velocity fields generated by krillbot using HOV kinematics with 0° (A-D), and 20° body angle (E-H) kinematics of *E. superba* [20]. (A & E) Start of power stroke (PS). (B & F) Middle of PS. (C & G) End of PS, which coincides with the start of recovery stroke (RS). (D & H) Middle of RS. All time points are in reference to P5.

In addition to changing the body angle of the krillbot, we also examined the effect of changing the phase lag between pleopods on the wake. **Figure 7** shows the wake of the krillbot (20° body angle) performing modified HOV kinematics with uniform *ϕ* of 0%, 25%, and 50% of cycle time at the same time points shown in **Figure 6**. Synchronous motion (0% phase lag, left) shows the wake splitting into two jets, with one directed downward and toward the tail, and the other directed downward and slightly forward. This forward wake would be expected to result in reduced swimming speed relative to a wake directed in the caudal direction only. 50% phase lag (right) shows a wake directed in a primarily downward direction. This wake is broader but slower than the wake generated by the unmodified HOV kinematics (**Figure 6**). An organism swimming in a quiescent fluid that generates a faster, narrower wake with the same mass flux is expected to generate higher thrust relative to a wider, slower wake. When this thrust vector is angled downward, this allows the propulsive force to be used for weight support, as in hovering krill. Unlike the cases of 0% and 50% phase lag, the 25% phase lag results in wake with a narrow, fast-moving jet directed in a primarily downward direction, which would provide an organism with the most thrust. This is consistent with both observed behaviors in krill, which use phase lags of approximately 15-30% during HOV [2, 20], and with computational results which suggest that phase lag of approximately 25% provides the largest average flux [35].

**Figure 7.**
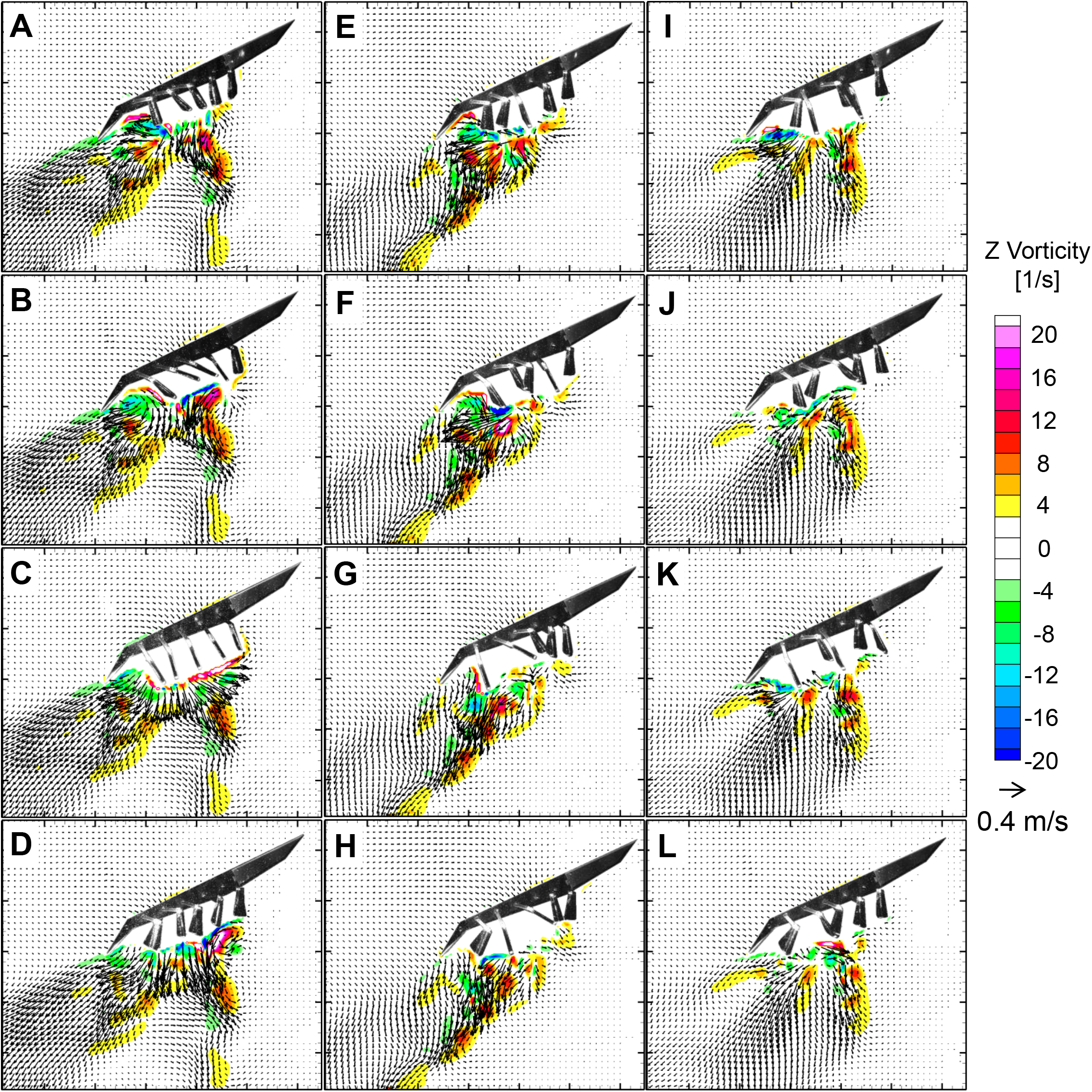
Out-of-plane vorticity contours overlaid with velocity fields generated by krillbot (20° body angle) using modified HOV kinematics of *E. superba* such that phase lag between adjacent pleopods were equal. (A-D) synchronous motion (0% phase lag), (E-H) 25% phase lag. (I-L) 50% phase lag. Time points for each row are the same as in **Figure 6**.

PIV was also performed for FFW kinematics, with unmodified (0 and 20° body angle) kinematics and for modified FFW kinematics with 0% and 15% phase lag at 20° body angle shown in **Supplementary Figures S4-S5**. At 0°, FFW motion generates two distinct jets, one in a downward direction, and the other in a primarily ventral direction. In contrast, the wake at 20° body angle does not have distinct jets (**Figure S4**). For changing phase lag, the *ϕ*=0% case shows a periodic motion while *ϕ*=15% shows a straighter, more continuous jet (**Figure S5**). However, the wakes here should not be taken as true indicators of the wake of *E. superba* using FFW gait, since the normalized speed and displacement efficiency of the krillbot was found to be much lower than krill (**Table 2**), due to drag forces on the support system that resist krillbot motion.

### 3.5 Comparison to swimming euphausiids

The krillbot is approximately 11 times larger than the *E. superba* individuals recorded by Murphy et al. [20], but is geometrically similar and operates at approximately the same Reynolds numbers and Strouhal numbers. Protopodite kinematics were prescribed to match data previously reported on freely-swimming *E. superba* [20] and matched relatively well in the krillbot (**Figures 3A-3B**). Normalized swimming speed, displacement efficiency, and advance ratio of the krillbot (**Table 2**) were lower than for the two species of freely-swimming krill [20,25]. Reynolds number for the krillbot was designed to match the Reynolds number range for Antarctic krill [20-21], and Strouhal numbers (dependent on momentum transfer to the wake) were matched as well. Though kinematics for this study were based on Antarctic krill, they could be easily modified to match other species as such data becomes available. Lower advance ratios are observed for the krillbot despite using kinematics identical to those of *E. superba*. This suggests that the relatively lower swimming performance of the krillbot is due primarily to added drag from the assembly (cabling resistance, drag generated by krillbot suspension), and possibly also due to restriction of one-dimensional travel. Additionally, the simplified geometries of the rigid krillbot pleopods, and the design of their hinges, could contribute to lower thrust coefficients in the krillbot than in live animals.

## 4 Discussion

Although several studies of metachronal propulsion have been performed in freely-swimming crustaceans [4,19,21] and using numerical [2,34-36] and robotic models [33,37], the relative effects of varying individual stroke kinematic parameters on free-swimming performance are unclear. In this study, we examined free-swimming performance as a function of inter-pleopod phase lag (*ϕ*) using a self-propelled biomimetic krill robot. Krillbot pleopods were programmed to move using previously published fast-forward swimming (FFW) and hovering (HOV) kinematics of freely-swimming Antarctic krill, *E. superba* [20]. Additionally, *ϕ* was varied from 0% of the stroke period to a maximum of 35% (FFW) or 50% (HOV). Swimming performance was assessed using free-swimming speed and tethered thrust. Regardless of phase lag, metachronal motion of pleopods (non-zero *ϕ*) resulted in increasing swimming speed and thrust compared to synchronous paddling (*ϕ*=0%). Flow visualization showed dramatic differences between different phase lags. The angled jet generated by HOV can enable downward momentum transfer for animal weight support needed during *E. superba* feeding. Further, the coherent wake structure generated by paddling with non-zero *ϕ* could potentially assist in hydrodynamic signaling between neighboring krill. To the best of the authors’ awareness, this is the first study to report: 1) thrust generated by FFW and HOV kinematics of *E. superba;* 2) effect of varying *ϕ* on free-swimming performance; 3) flow generated by HOV kinematics of *E. superba* with changing body angle and phase lag; and 4) on the fluid dynamic mechanism underlying thrust augmentation by metachronal rowing (i.e., non-zero *ϕ*).

FFW kinematics with phase lags (*ϕ*) ranging between 15% to 25%, similar to the range of *ϕ* reported in FFW motion of freely-swimming *E. superba*, were found to provide highest thrust and swimming speed. This is also within the range of *ϕ* where highest average volumetric flux was obtained in previous computational modeling studies of tethered metachronal propulsion [35-36], as well as within the range of *ϕ* where largest fluid momentum was observed in a previous study using a tethered metachronal robotic model [33]. Zhang et al. [35] argued that geometries generated by adjacent pleopods stroking at *ϕ*=25% could help in entrapping more volume of fluid in the inter-pleopod gap during PS and less volume of fluid during RS. However, they did not examine free-swimming performance with their modeling approach.

Despite simplifications in the krillbot pleopod design, including the lack of ‘lobed’ structure (with endopodite and exopodite), rigid structure, and the absence of setae, our robotic model characteristics compare well with those of free-swimming *E. superba* and *E. pacifica* (**Table 2**). The krillbot can reasonably mimic the pleopod kinematics associated with two distinct *E. superba* swimming gaits (FFW and HOV). Strouhal number of the krillbot matches that of *E. superba* when the geometry, Reynolds number, and kinematics are the same. However, losses in the system (resistance offered by cabling, drag generated by suspension system) inevitably result in lower displacement efficiency in the krillbot (particularly when using FFW kinematics) than in euphausiids.

Changing the body angle of the krillbot from 0° to 20° resulted in lower swimming speeds. Two results of changing the body angle are the body drag being greater for the same swimming speed at 20° due to larger cross-sectional area, as well as the momentum of the fluid wake generated by the paddling motion being directed in a more downward direction. These two factors contribute to the lower swimming speed of the krillbot when positioned with a 20° body angle, as compared to a 0° body angle, and help paddling krill maintain their position while still supporting their weight while swimming at reported body angles of 20-30° [20].

The flow generated by the krillbot using HOV kinematics was qualitatively similar to that reported in hovering *E. superba* [21], and the wake structure was found to change along with swimming performance for changing phase lag. The ventrally angled jet generated by HOV motion can enable downward momentum transfer for weight support needed during *E. superba* feeding [21]. Additionally, since increasing the body angle from 0° to 20° resulted in decreased swimming speed and a more vertical orientation of the wake, farther increasing the body angle could force the robot to paddle in position, which would be useful for hovering robots using this mechanism. Crustaceans are observed to change the angle of their tail relative to their bodies, and to change the angle of their abdomen relative to their thorax in order to achieve hovering and different maneuvers. Further studies are needed to identify the specific roles of body and tail angles, as well as other geometric and kinematic parameters.

To examine the fluid dynamic mechanism augmenting krillbot thrust for *ϕ*>0%, we characterized the change in distance between hinges of adjacent pleopods in time (**Figures 8A-8B**). For synchronous pleopod motion (*ϕ*=0%), the distance between hinges is nearly constant (≈42 mm, **Figure 8A**). However, adjacent pleopods move relative to each other when operating with non-zero phase lag (**Figure 8B**). By comparing the change in hinge distance to the krillbot displacement, we found that the krillbot motion slows when adjacent pleopods move towards each other and speeds up when adjacent pleopods move apart (**Figures 8C-8D**). This points to a mechanism of thrust generation whereby strong tip vortices generated during the power stroke [43] create suction to pull the krillbot forward, rather than a mechanism by which the expulsion of jets between paddles pushes the krillbot. While jets will be expelled from between adjacent paddles moving towards each other with a phase lag, they will be directed in a primarily vertical direction, particularly when there is a non-zero body angle. Jointed pleopods help to direct the jet flow downward when pleopods move towards each other, in contrast to the primarily horizontal wake generated when two adjacent paddles move apart from each other.

**Figure 8.**
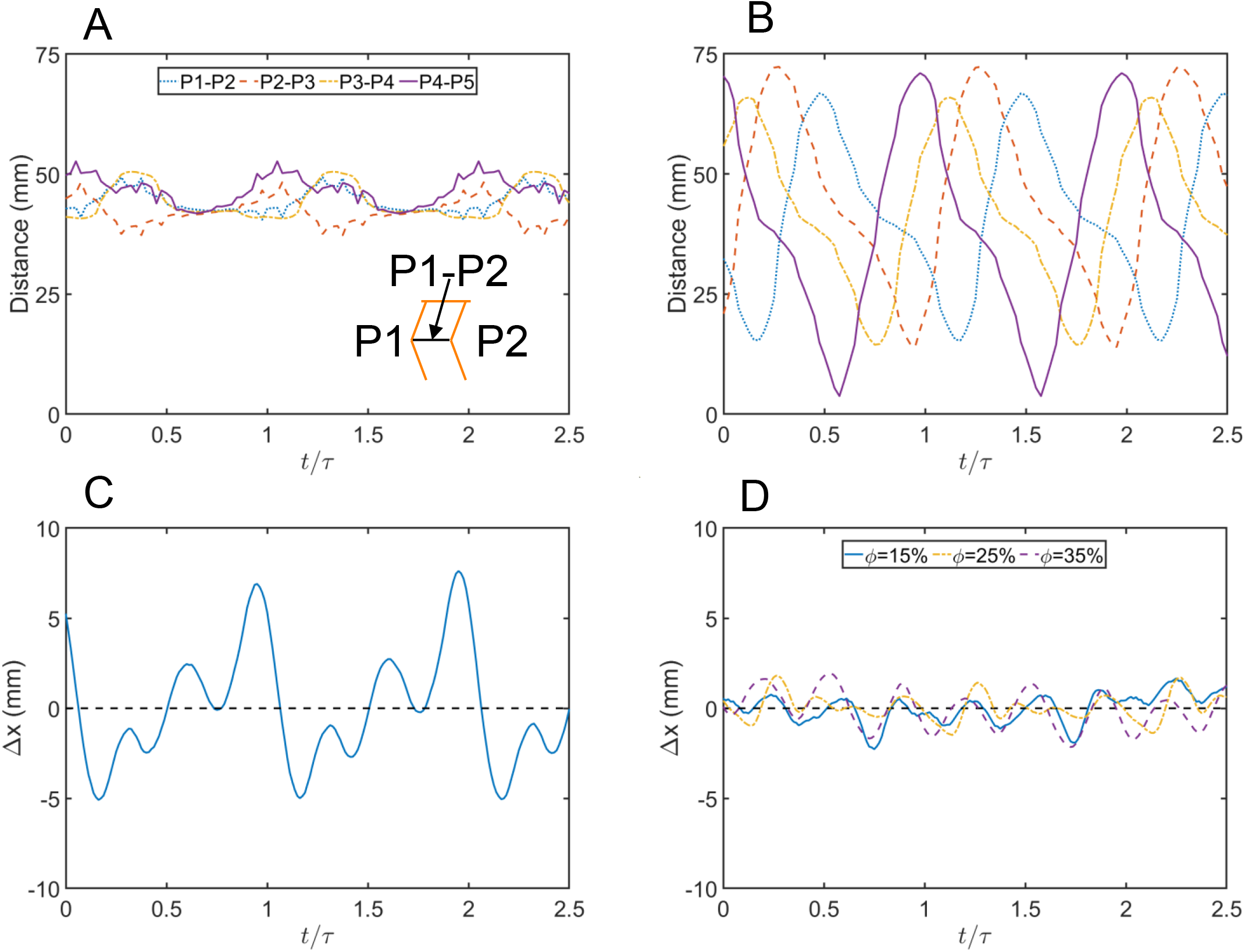
Inter-hinge distances and body position deviation (from mean swimming speed) for FFW kinematics with uniform *ϕ*. Body position deviation (*Δx*) was calculated using equation (8). Inter-hinge distance between pleopod pairs for FFW kinematics was nearly constant at 0% phase lag (A) as compared to periodic fluctuations for 15% phase lag (B). Variation in *Δx* was larger for *ϕ*=0% (shown in C) as compared to *ϕ*=15%-35% (shown in D).

## 5 Conclusions

Using a self-propelling krill robot, we show that the phase lag range used by freely-swimming Antarctic krill (*E. superba*) is well-suited for achieving peak swimming performance (FFW), and for downward transfer of momentum (HOV). Variation of inter-pleopod phase lag in metachronal swimming directly affects steady swimming speed during self-propulsion. Swimming speed increases at the start of each paddle’s power stroke, when the tip vortices grow the fastest [43]. Swimming speed decreases when the paddles move closer together. Collectively, these findings illustrate the importance of phase lag in free-swimming performance of coordinated appendage rowing and clear the way for future studies to examine the effects of other morphological and kinematic parameters on swimming performance.

## Ethics

This article does not present research with ethical considerations.

## Data Accessibility Statement

Electronic supplementary material consisting of representative high-speed videos of krillbot motion, and numerical data used in figures 3, 6, 7 and 8 are available within Figshare: https://figshare.com/s/030703a893e43ef4cfa8 [44]. (Note: this is a private link for the review process and will be made public with a DOI if/when manuscript is accepted).

## Authors’ Contributions

M.P.F. and A.S. conceived of and designed the study. M.P.F. acquired experimental data. M.P.F. and A.S. analyzed the data, drafted and critically revised the manuscript. All authors gave final approval for publication and agree to be held accountable for the work performed therein.

## Competing Interests

We declare we have no competing interests.

## Acknowledgements

We would like to thank Tyler Blackshare for his assistance in digitizing the krillbot pleopod motion.

## Funding

This work was supported by the National Science Foundation (grant number CBET 1706762 to A.S.).

## Notes

### Competing Interest Statement

The authors have declared no competing interest.

### Summary of Updates

Revision following first round of peer-review.

## References

1. Kils U. 1981 Swimming behaviour, swimming performance and energy balance of Antarctic krill Euphausia superba. BIOMASS Sci. Ser. 3, 1–21.

2. Alben S, Spears K, Garth S, Murphy D, Yen J 2010 Coordination of multiple appendages in drag-based swimming J. R. Soc. Interface 7, 1545–1557

3. Lenz P H, Takagi D, Hartline D K 2015 Choreographed swimming of copepod nauplii J. R. Soc. Interface 12

4. Campos E O, Vilhena D, Caldwell R L 2012 Pleopod rowing is used to achieve high forward swimming speeds during the escape response of Odontodactylus havanensis (Stomatopoda) J. Crustacean Biol. 32, 171–179

5. Van Duren L A, Videler J J 2003 Escape from viscosity: The kinematics and hydrodynamics of copepod foraging and escape swimming J. Exp. Biol. 206, 269–279

6. Sleigh, M A, & Barlow, D I 1980 Metachronism and control of locomotion in animals with many propulsive structures. In: Elder, H Y and Trueman, E R, (ed) Aspects of Animal Movement, Cambridge University Press, Cambridge, pp 49–70

7. Larson M, Kiger K T, Abdelaziz K, Balaras E 2014 Effect of metachronal phasing on the pumping efficiency of oscillating plate arrays Exp. Fluids 55

8. Sensenig A T, Kiger K T, Shultz J W 2009 The rowing-to-flapping transition: Ontogenetic changes in gill-plate kinematics in the nymphal mayfly Centroptilum triangulifer (Ephemeroptera, Baetidae) Biol. J. Linn. Soc. 98, 540–555

9. Sensenig A T, Kiger K T, Shultz J W 2010 Hydrodynamic pumping by serial gill arrays in the mayfly nymph Centroptilum triangulifer J. Exp. Biol. 213, 3319–3331

10. Sleigh M A, Blake J R, Liron N 1988 The propulsion of mucus by cilia Am. Rev. Respir. Dis. 137, 726–741

11. Wong L B, Miller I F, Yeates D B 1993 Nature of the mammalian ciliary metachronal wave J. Appl. Physiol. 75, 458–467

12. Elgeti J, Gompper G 2013 Emergence of metachronal waves in cilia arrays Proc. Natl. Acad. Sci. U. S. A. 110, 4470–4475

13. Chateau S, Favier J, Poncet S, D’Ortona U 2019 Why antiplectic metachronal cilia waves are optimal to transport bronchial mucus Phys. Rev. E 100, 42405

14. Bandyopadhyay P R, Hansen J C 2013 Breakup and then makeup: A predictive model of how cilia self-regulate hardness for posture control Sci. Rep. 3, 1–10

15. Jana S, Um S H, Jung S 2012 Paramecium swimming in Capillary Tube Phys. Fluids 041901

16. Funfak A, Fisch C, Abdel Motaal H T, Diener J, Combettes L, Baroud C N, Dupuis-Williams P 2015 Paramecium swimming and ciliary beating patterns: A study on four RNA interference mutations Integr. Biol. (United Kingdom) 7, 90–100

17. Catton K B, Webster D R, Brown J, Yen J 2007 Quantitative analysis of tethered and free-swimming copepodid flow fields J. Exp. Biol. 210, 299–310

18. Jiang H, Kiørboe T 2011 The fluid dynamics of swimming by jumping in copepods J. R. Soc. Interface 8, 1090–1103

19. Schabes M and Hamner W 1992 Mysid locomotion and feeding: kinematics and water-flow patterns of Antarctomysis sp., Acanthomysis sculpta, and Neomysis rayii J. Crustacean Biol. 12, 1–10

20. Murphy D W, Webster D R, Kawaguchi S, King R, Yen J 2011 Metachronal swimming in Antarctic krill: Gait kinematics and system design Mar. Biol. 158, 2541–2554

21. Murphy D W, Webster D R, Yen J 2013 The hydrodynamics of hovering in Antarctic krill Limnol. Oceanogr. Fluids Environ. 3, 240–255

22. Lim J L, DeMont M E 2009 Kinematics, hydrodynamics and force production of pleopods suggest jet-assisted walking in the American lobster (Homarus americanus) J. Exp. Biol. 212, 2731–2745

23. Wiese K, Ebina Y 1995 The propulsion jet of Euphausia superba (Antarctic Krill) as a potential communication signal among conspecifics J. Mar. Biol. Assoc. United Kingdom 75, 43–54

24. Zhou M, Dorland R D 2004 Aggregation and vertical migration behavior of Euphausia superba Deep. Res. Part II Top. Stud. Oceanogr. 51, 2119–2137

25. Catton K B, Webster D R, Kawaguchi S, Yen J 2011 The hydrodynamic disturbances of two species of krill: Implications for aggregation structure J. Exp. Biol. 214, 1845–1856

26. Murphy D W, Olsen D, Kanagawa M, King R, Kawaguchi S, Osborn J, Webster D R, Yen J 2019 The three dimensional spatial structure of Antarctic krill schools in the laboratory Sci. Rep. 9, 1–12

27. Atkinson A, Siegel V, Pakhomov E A, Jessopp M J, Loeb V 2009 A re-appraisal of the total biomass and annual production of Antarctic krill Deep. Res. Part I Oceanogr. Res. Pap. 56, 727–740

28. Swadling K M, Ritz D A, Nicol S, Osborn J E, Gurney L J 2005 Respiration rate and cost of swimming for Antarctic krill, Euphausia superba, in large groups in the laboratory Mar. Biol. 146, 1169–1175

29. Kawaguchi S, Kilpatrick R, Roberts L, King, RA, Nicol, S 2011 Ocean-bottom krill sex. Journal of plankton research, 33(7), 11341138.

30. Endo Y 1993 Orientation of Antarctic Krill in an Aquarium Nippon Suisan Gakkaishi 59, 465–468

31. Marschall H P 1988 The overwintering strategy of Antarctic krill under the pack-ice of the Weddell Sea Polar Biol. 9, 129–135

32. Hamner W M 1984 Aspects of schooling in Euphausia superba J. Crustacean Biol. 4, 67–74

33. Ford M P, Lai H K, Samaee M, Santhanakrishnan A 2019 Hydrodynamics of metachronal paddling: effects of varying Reynolds number and phase lag R. Soc. Open Sci. 6, 191387

34. Takagi D 2015 Swimming with stiff legs at low Reynolds number Phys. Rev. E 92, 023020

35. Zhang C, Guy R D, Mulloney B, Zhang Q, Lewis T J 2014 Neural mechanism of optimal limb coordination in crustacean swimming Proc. Natl. Acad. Sci. U. S. A. 111, 13840–13845

36. Granzier-Nakajima S, Guy R D, Zhang-Molina C 2020 A Numerical Study of Metachronal Propulsion at Low to Intermediate Reynolds Numbers Fluids 5, 86

37. Hayashi R, Takagi D 2020 Metachronal swimming with rigid arms near boundaries Fluids 5, 24

38. Kils, U 2011 “Antarctic krill (Euphausia superba).jpg”, used under Creative Commons Attribution-Share Alike 3.0 Unported license / Background removed and dimensional annotations added

39. Schneider C A, Rasband W S, Eliceiri K W 2012 NIH Image to ImageJ: 25 years of image analysis Nat. Methods 9, 671–675

40. Taylor G K, Nudds R L, Thomas A L R 2003 Flying and swimming animals cruise at a Strouhal number tuned for high power efficiency Nature 425, 705–707

41. Hedrick T L 2008 Software techniques for two-and three-dimensional kinematic measurements of biological and biomimetic systems Bioinspiration and Biomimetics 3

42. Yen J, Brown J, Webster D R 2003 Analysis of the flow field of the krill, Euphausia pacifica Mar. Freshw. Behav. Physiol. 36, 307–319

43. Kim D, Gharib M 2011 Characteristics of vortex formation and thrust performance in drag-based paddling propulsion. J. Exp. Biol. 214(13), 2283-2291.

44. Ford M, Santhanakrishnan A 2020 Data from: On the role of phase lag in multi-appendage metachronal swimming of Euphausiids. Figshare private link (will be made public with a DOI if/when manuscript is accepted): https://figshare.com/s/030703a893e43ef4cfa8.

