## Supplementary Figures 1 to 5 for "On the role of phase lag in multi-appendage metachronal swimming of euphausiids"

**Figure S1.** Time series of velocity for modified FFW kinematics at 0º body angle (left) and 20º body angle (right). Each row represents a different uniform phase lag, 0%, 15%, 25%, 35% from top to bottom. Blue lines represent instantaneous velocity, while red lines represent the average velocity over one cycle ($\pm\frac{1}{2}$ cycle form the time point indicated). A negative instantaneous value means the krillbot is moving backward, while a positive value means it is moving forward. FFW at 0º body angle is generally faster than at 20º, and 0% phase lag has the most velocity fluctuation, while 15% has the least.

**Figure S2.** Time series of velocity for modified HOV kinematics at 0º body angle (left) and 20º body angle (right). Each row represents a different uniform phase lag, 0%, 15%, 25%, 35%, 50% from top to bottom. Blue lines represent instantaneous velocity, while red lines represent the average velocity over one cycle ($\pm\frac{1}{2}$ cycle form the time point indicated).

**Figure S3.** Time series of velocity for unmodified kinematics at 0º body angle (top) and 20º body angle (bottom). Left is FFW kinematics, while right is HOV kinematics. Blue lines represent instantaneous velocity, while red lines represent the average velocity over one cycle ($\pm\frac{1}{2}$ cycle form the time point indicated).

**Figure S4.** Out-of-plane vorticity contours overlaid with velocity fields generated by krillbot using unmodified FFW kinematics with 0º (A-D), and 20º body angle (E-H). (A & E) Start of power stroke for P5 (PS). (B & F) Middle of PS. (C & G) End of PS, which coincides with the start of recovery stroke (RS). (D & H) Middle of RS.

**Figure S5.** Out-of-plane vorticity contours overlaid with velocity fields generated by krillbot using modified FFW kinematics with and 20º body angle. (A-D) show synchronous motion while (E-H) show 15% phase lag. (A & E) Start of power stroke for P5 (PS). (B & F) Middle of PS. (C & G) End of PS, which coincides with the start of recovery stroke (RS). (D & H) Middle of RS.

### Figure S1

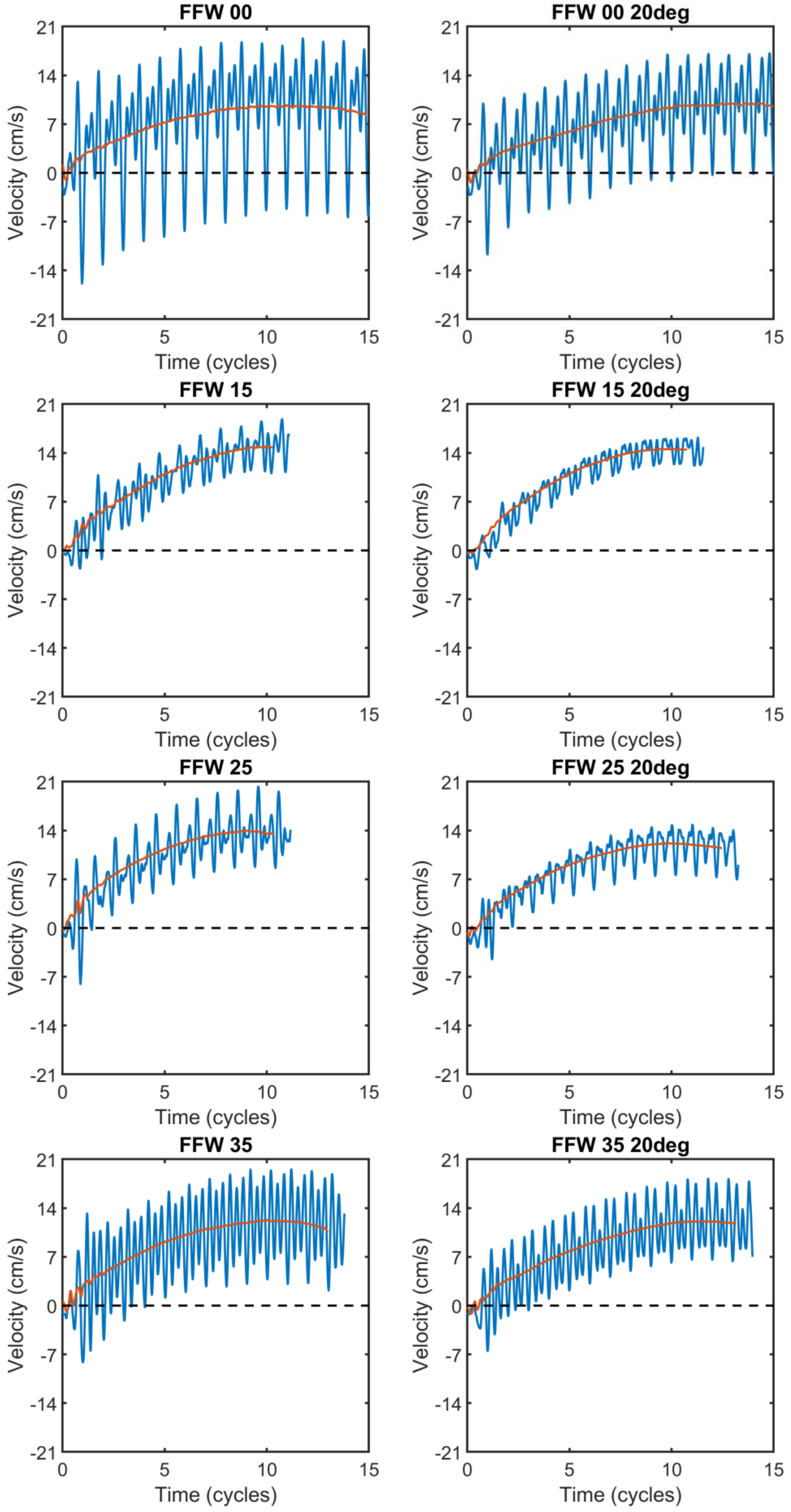

FFW 00
FFW 00 20deg
FFW 15
FFW 15 20deg
FFW 25
FFW 25 20deg
FFW 35
FFW 35 20deg
Velocity (cm/s)
Time (cycles)
21
14
7
0
-7
-14
-21
0
5
10
15

Figure S2

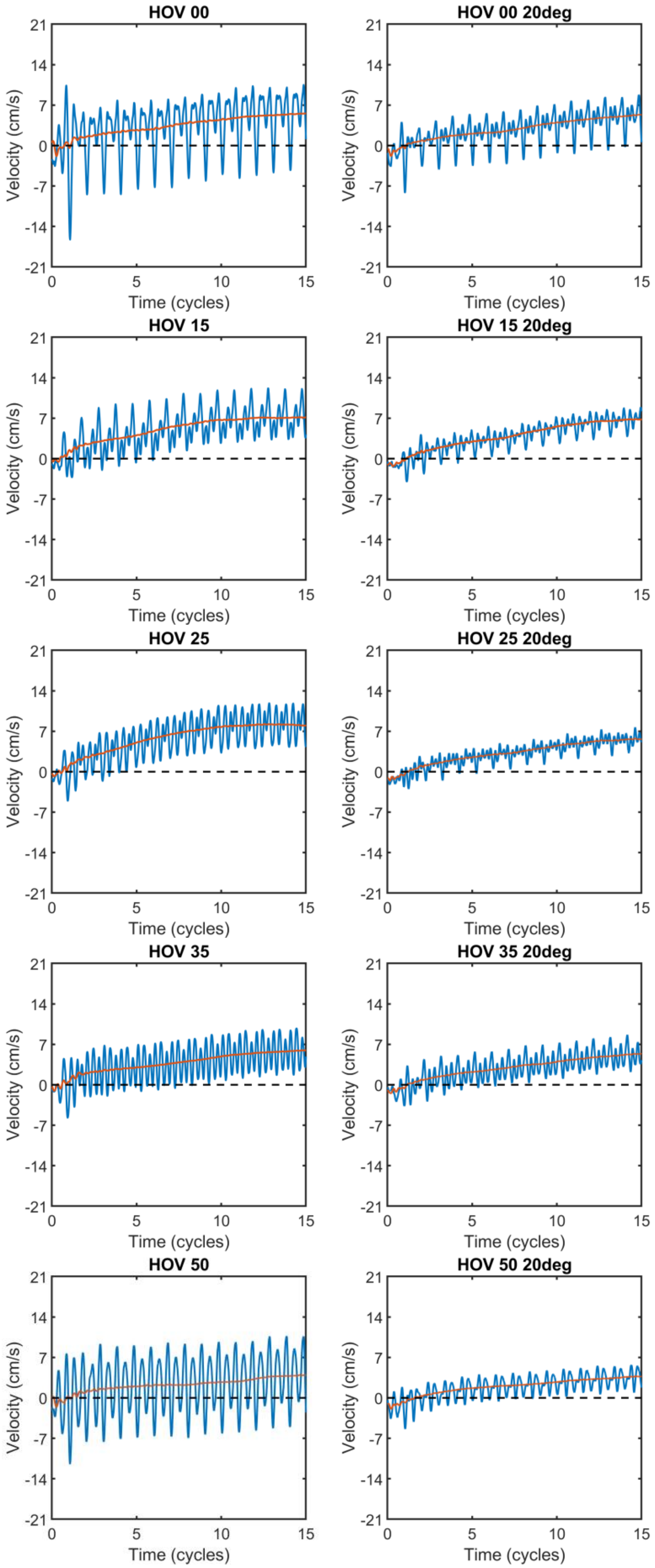

HOV 00
HOV 00 20deg
HOV 15
HOV 15 20deg
HOV 25
HOV 25 20deg
HOV 35
HOV 35 20deg
HOV 50
HOV 50 20deg
Velocity (cm/s)
Time (cycles)
21
14
7
0
-7
-14
-21
0
5
10
15

Figure S3

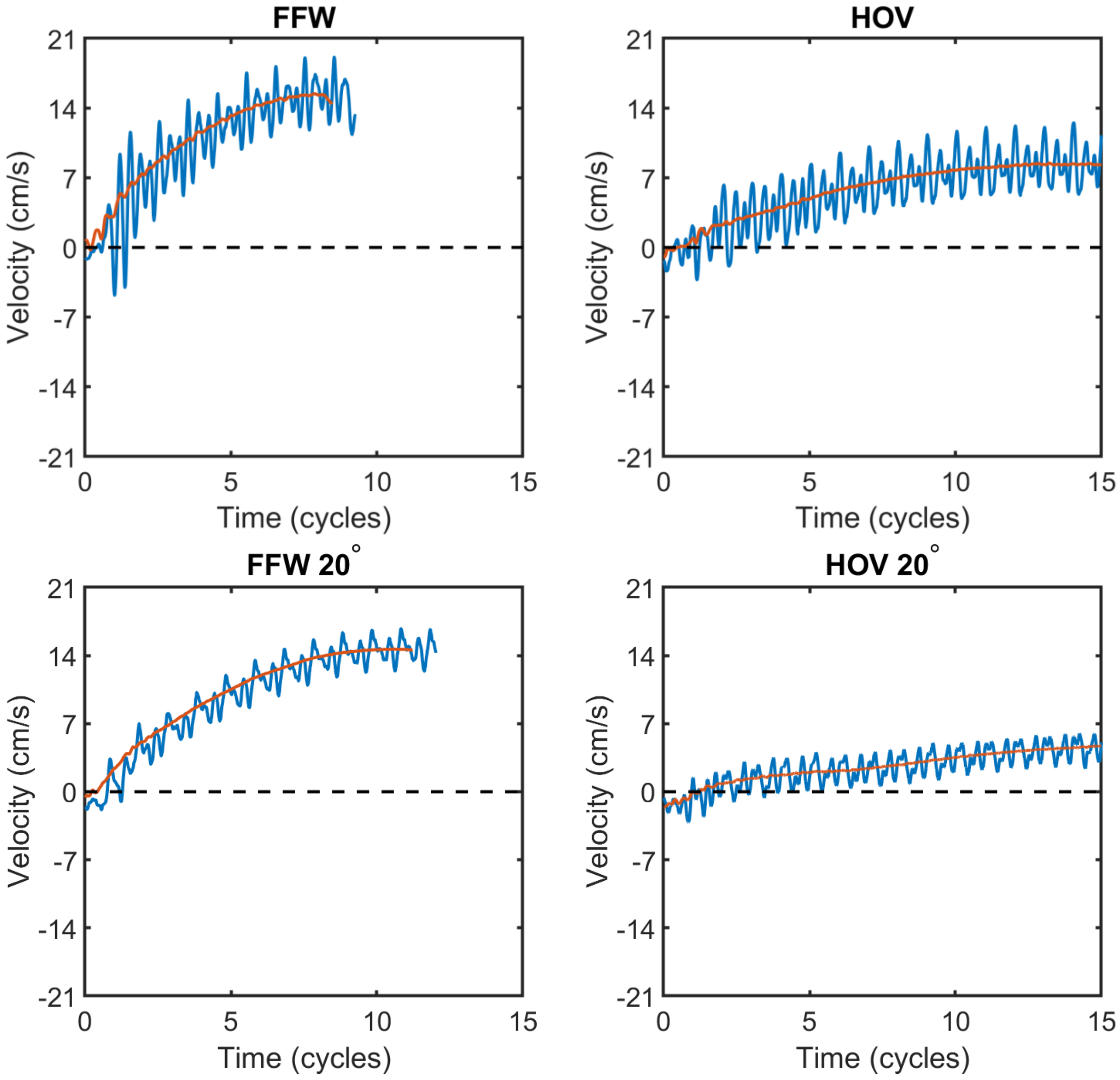

FFW
HOV
FFW 20°
HOV 20°
Velocity (cm/s)
Time (cycles)
21
14
7
0
-7
-14
-21
0
5
10
15

### Figure S4

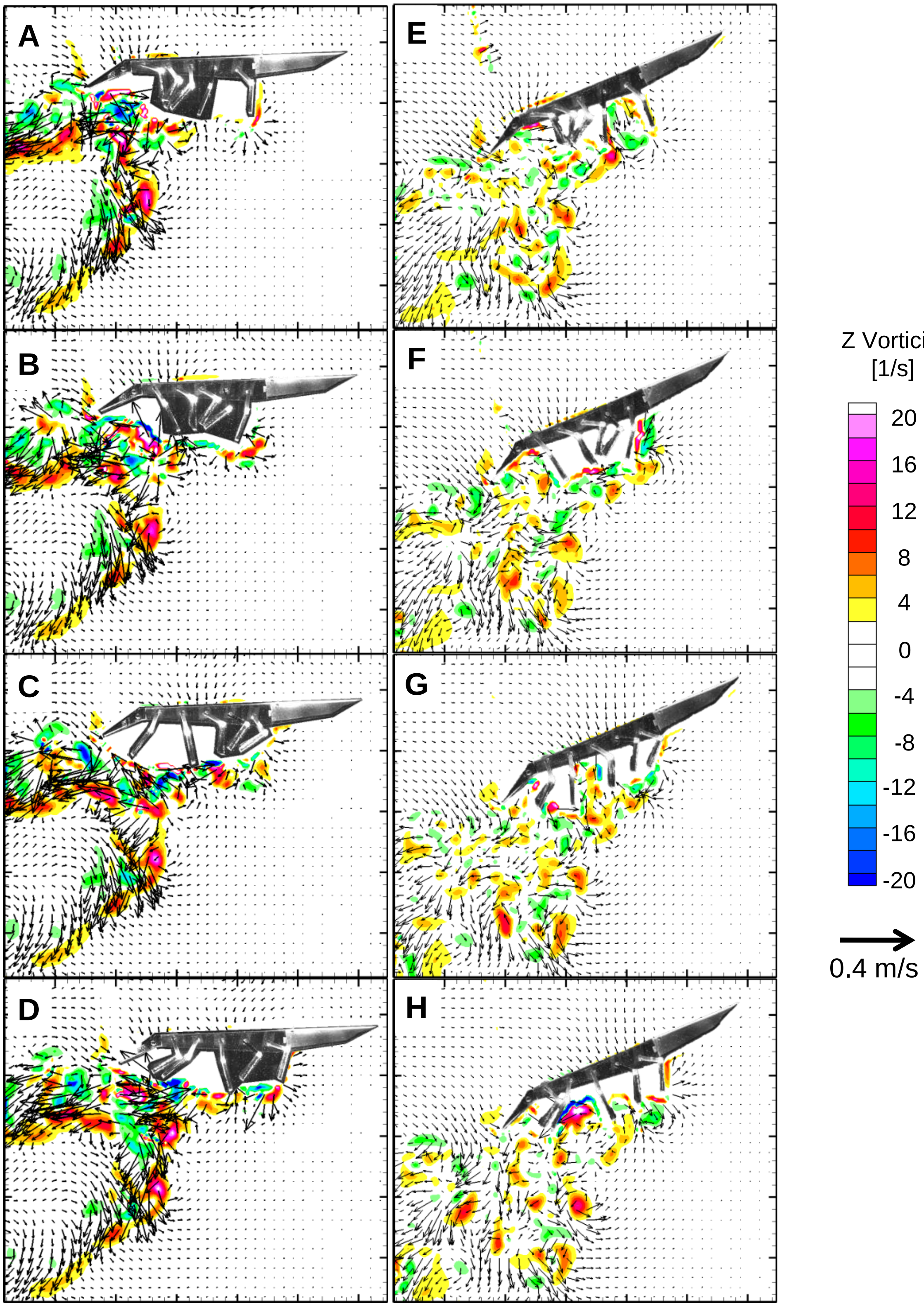

Figure S5

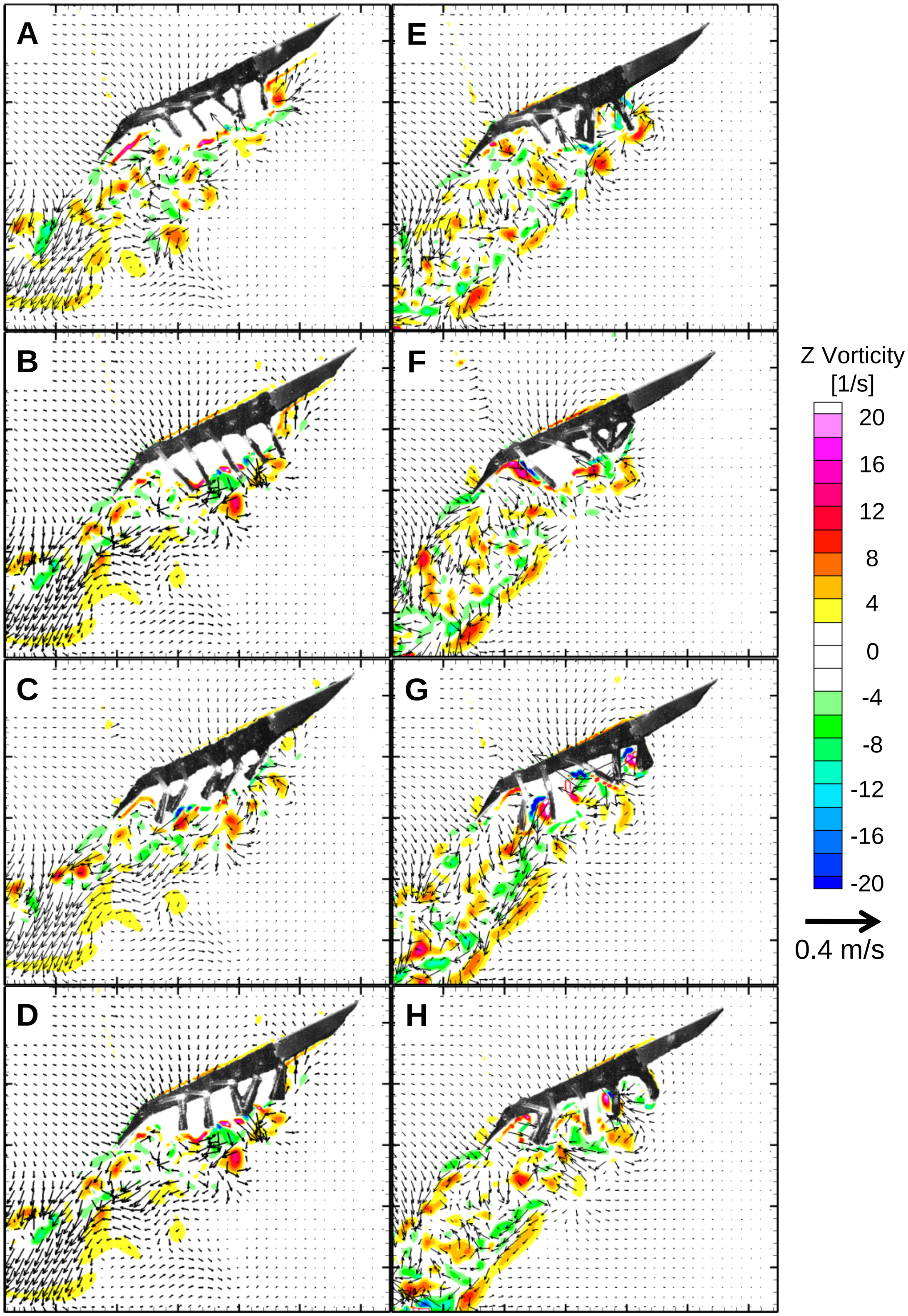
